# A Sterol-PI(4)P exchanger controls the Tel1/ATM axis of the DNA Damage Response

**DOI:** 10.1101/2022.07.13.499867

**Authors:** Sara Ovejero, Sylvain Kumanski, Caroline Soulet, Julie Azarli, Benjamin Pardo, Olivier Santt, Angelos Constantinou, Philippe Pasero, María Moriel-Carretero

## Abstract

Upon DNA damage, cells activate the DNA Damage Response (DDR) to coordinate proliferation and DNA repair. Dietary, metabolic, and environmental inputs are emerging as modulators of how DNA surveillance and repair take place. Lipids hold potential to convey these cues, although little is known about how. We observed that lipid droplet (LD) number specifically increased in response to DNA breaks. We show that the selective storage of sterols into these LD concomitantly stabilizes phosphatidyl-4-inositol (PI(4)P) at the Golgi, where it binds the DDR kinase ATM. In turn, this titration attenuates the initial nuclear ATM-driven response to DNA breaks, thus allowing processive repair. Further, manipulating this loop impacts the kinetics of DNA damage signaling and repair in a predictable manner. Thus, our findings have major implications for tackling genetic instability pathologies through dietary and pharmacological interventions.

## Introduction

The integrity of the genetic information needs to be preserved to warrant cell and organism fitness. The cell therefore possesses surveillance and repair strategies to accomplish this task. Cells challenged with DNA damage or other sources of genotoxic stress activate the DNA Damage Response (DDR) to arrest the cell cycle in order for DNA repair to take place. This occurs upon lesion detection by dedicated sensors, which engage the upstream kinases of this response, namely Tel1/ATM (as known in *S. cerevisiae* and humans, respectively) and Mec1/ATR. Kinase activity promotes effector phosphorylation, among which Rad53/CHK1-CHK2, which in turn drive cell protection and recruit repair factors. Successful repair coupled to subsequent checkpoint inactivation is named recovery. However, if the damage cannot be repaired, different scenarios may emerge: cells can remain in a permanent cell-cycle arrest, which, from a developmental point of view, compromises future lineages. Alternatively, an exacerbated activation of the DDR triggers apoptosis, leading to cell death and tissue loss in many degenerative diseases ^1, 2^. Last, cells could resume cell cycle in the presence of the damage. This latter phenomenon is called adaptation and permits cell survival while propagating genome instability. Adaptation is achieved by downregulation of the DDR and is reported as a cell survival strategy under chronic DNA damage conditions in unicellular organisms ^3, 4^. In multicellular organisms, adaptation is generally prevented through the engagement of apoptosis or senescence, which limit unrestrained proliferation ^5^. Overall, the intensity and the context of the DDR activation dictates repair, apoptosis or uncontrolled proliferation.

Another important aspect ruling cell fate relies on lipids. This is clearly illustrated during cancer development, in which lipids act as energy sources and signaling nodes, promoting progression and even metastasis ^6, 7^. Saturated fatty acids also make cells less permeable to exogenous toxins, oxidative damage and chemotherapeutic agents ^8^. A specific player in the metabolism of lipids are lipid droplets (LD), the sole organelle to be delimited by a monolayer of phospholipids. LD are classically known as the cell storage for fats and were long seen as a static, almost inert bag of lipids. They are mainly filled with apolar lipids such as esterified fatty acids in the form of triacylglycerols (TAGs) and esterified sterols in the shape of steryl esters (STEs). They are born from the Endoplasmic Reticulum (ER) into the cytoplasm, where their half-life is regulated by an intricate interplay of esterifying enzymes, LD-ER contact factors and lipases ^9^. Their relevance is illustrated by the wide spectrum of diseases that derive from their mishandling ^10^. Yet, even if in cancer the presence of LD is a hallmark of bad prognosis, relapse and resistance to chemotherapy ^11–13^, the specific relevance of LD in cell fate decisions is less understood. Last, a further important aspect of lipid regulation occurs at membrane contact sites. These are locations where membranes from two (or more) different organelles approach, being no longer than 80 nm apart, and where specific protein machines execute the transfer of lipids from one membrane to another, creating fluxes of functional importance^14^. Among them, the interface between the Trans-Golgi and the ER hosts the OSBP1 exchanger: sterol moieties present in the ER are extracted from this location and inserted in the Trans-Golgi in exchange of phosphatidyl-inositol-4-phosphoate (PI(4)P) molecules, which are in turn extracted from the Golgi and inserted in the ER. Once in the ER, these PI(4)P molecules are fast hydrolyzed by the phosphatase SAC1, which boosts the activity of this lipid exchange machinery ^15^. Thus, the low sterol content of the ER is warranted by the fine regulation of the activity of OSBP1.

Observations from the last seven years suggest links between nutrient sensing, lipidic programs and DNA damage handling ^16–21^, yet a deep mechanistic understanding is still missing. In this work, we explore whether the response of the cell to DNA damage can be modulated by the metabolism of lipids. We found that human ATM and *S. cerevisiae* Tel1, upstream kinases of the DDR and ancestral phosphatidyl-inositol kinases ^22^, can bind PI(4)P. In response to DSBs, the formation of LD that specifically store sterols is promoted, thus decreasing sterol levels at the ER. In turn, this limits the extraction of PI(4)P from the Golgi. The PI(4)P moieties stabilized this way keep ATM/Tel1 locked away from the nucleus, a titration that is key to permit efficient DNA repair. Last, we provide proof-of-concept that the manipulation of sterol metabolism at the interface between the Trans-Golgi and the ER permits an unanticipated control of ATM/Tel1, for example in the dampening of the DDR. Our data support the existence of an evolutionarily conserved mechanism whose control in response to DSBs can be tailored genetically and pharmacologically by manipulating the metabolism of lipids, thus bearing an exciting potential both in basic and applied research on genome stability.

## Results

### Lipid Droplets accumulate in response to DNA double strand breaks

To determine whether genotoxic stress impacts the accumulation of LD in cells, we performed kinetic studies by exposing asynchronous cultures of wild type (WT) *Saccharomyces cerevisiae* cells to genotoxic agents that induce DNA breakage (Figure 1A,B). We used zeocin, a radiomimetic agent that creates DNA single-stranded and double-stranded DNA breaks (DSBs) and camptothecin (CPT), which covalently traps topoisomerase I to DNA and leads to the accumulation of DSBs during DNA replication. DSB induction with zeocin or CPT triggered an increase of Rad52 foci formation, which promotes DSB repair by homologous recombination (HR) (Figure S1A) ^23^. The presence of DSBs is expected to activate the DNA damage response (DDR), which drives the accumulation of cells in the G_2_ phase of the cell cycle (2C DNA content in Figure 1B,C) ^24^. We then monitored the presence of LD using the vital dye Nile Red and observed that DSB-causing agents triggered a significant increase of LD in the cell population (Figure 1A-C & Figure S1C). We also exposed the cells to the genotoxic agent hydroxyurea (HU), which does not induce DSBs but blocks DNA replication by decreasing dNTP pools. Short time exposure to HU blocks cells in S phase (Figure 1D, right) and activates the DDR, but does not lead to the formation of Rad52 foci (Figure S1B) ^25^. In response to HU, LD did not accumulate in a significant manner (Figure 1D, left & Figure S1C), suggesting that they specifically accumulate in response to DSBs. We excluded that LD increase related to cells accumulation in G_2_, because treatment with nocodazole, which forces G_2_/M cell arrest (Figure 1E, right) due to microtubule depolymerization without creating DNA damage, failed to induce LD accumulation to the same extent (Figure 1E, left & Figure S1C). We therefore conclude that LD formation is stimulated in the event of DSBs.

**Figure 1.**
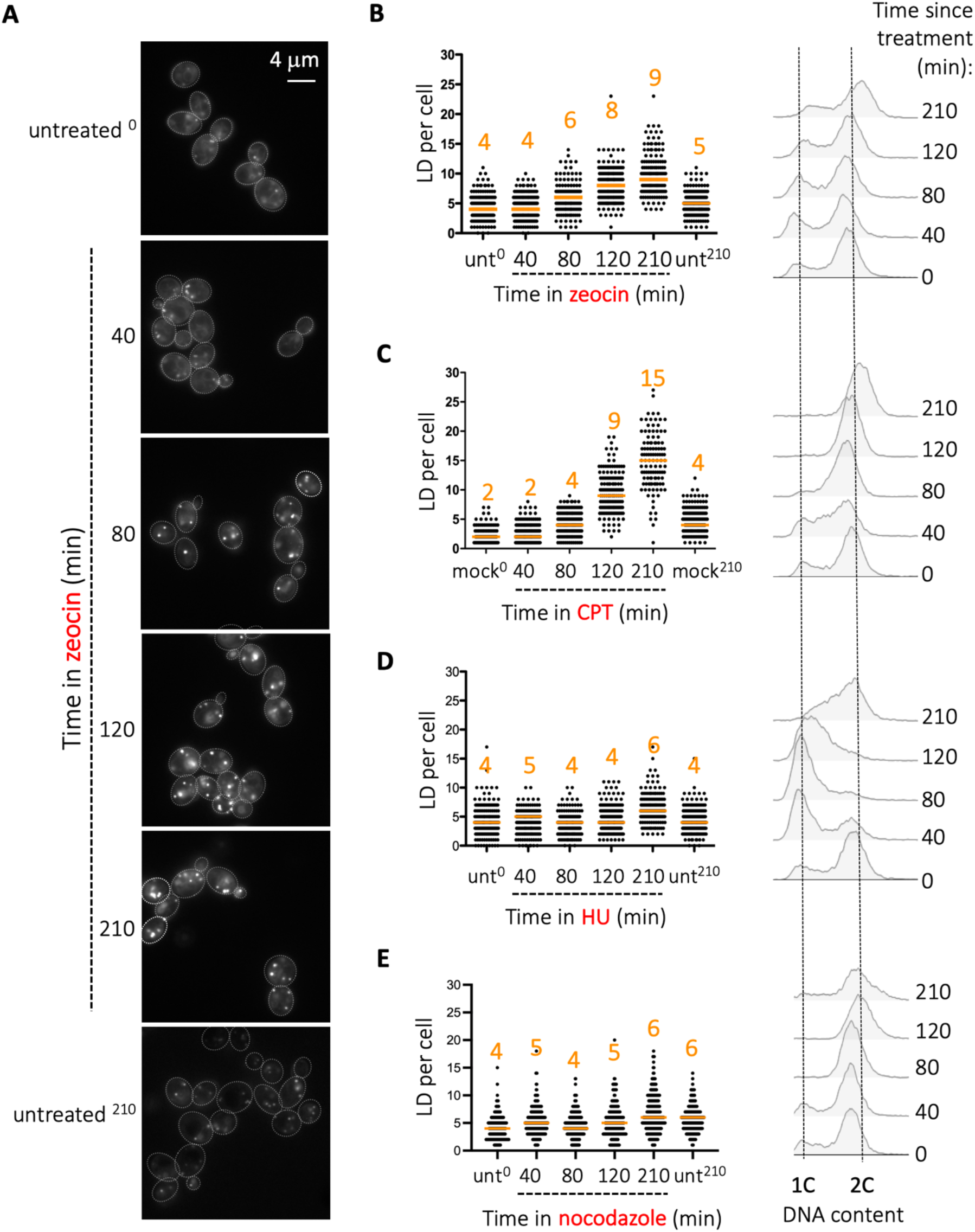
Ability of different DNA-damaging agents to induce lipid droplet accumulation. **A.** *S. cerevisiae* cells growing exponentially were treated (or not) with 100 µg/mL zeocin for the indicated times and stained using the vital dye Nile Red just prior to imaging to visualize LD. **B-E. Left:** At least 150 cells were counted per time point and the amount of LD per cell was inspected visually and plotted. The median value of each timepoint is indicated by an orange bar and number. Each panel corresponds to a representative time-course experiment in response to each indicated DNA-damaging agent (out of at least 3 per agent). “unt” = untreated condition. “mock” refers to the culture being treated with DMSO, the dissolving agent for CPT. Used concentrations were 100 µg/mL zeocin, 100 µM CPT, 100 mM HU and 15 µg/mL nocodazole. **Right:** the same cultures were analysed by cytometry to assess DNA content (1C = unreplicated DNA, 2C = fully replicated DNA) during the time-course to confirm that the used agents were causing cell arrest at the expected phases of the cell cycle.

### Inability to esterify sterols exacerbates DNA double strand break signaling

To assess whether the accumulation of LD in response to DSBs was of functional relevance, and to ascertain which type of LD contents could be implicated, we explored the sensitivity to DSB-making agents of cells in which the storage of different LD constituents was compromised. LD store two different types of apolar lipids, namely triacylglycerol (TAGs) and steryl esters (STEs), deriving from the esterification of fatty acids and sterols, respectively ^9^. We used isogenic strains lacking TAGs (*lro1Δ dga1Δ*, designated *tagΔ* hereafter) or STEs (*are1Δ are2Δ*, designated *steΔ* hereafter) ^26^. As expected, *tagΔ* and *steΔ* cells have a low basal level of LD (Figure S1D, upper panel and ^26^). We observed that only *steΔ* cells were more sensitive to zeocin and CPT than WT cells, yet not to HU (Figure 2A). This suggests that only the esterification of sterols into steryl esters then their storage in LDs is important to cope with the induction of DSBs.

**Figure 2.**
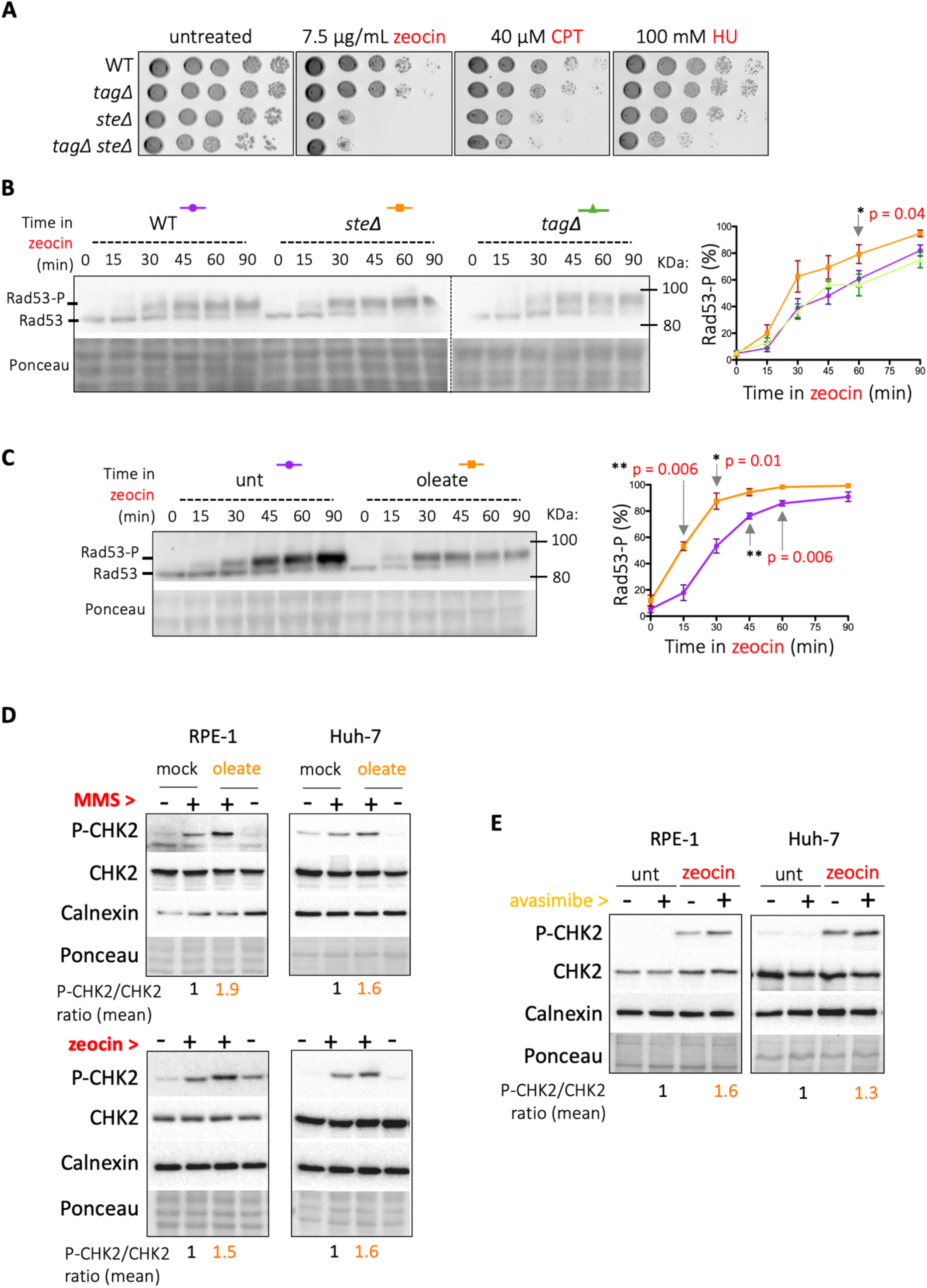
Lack of sterol storage or esterification exacerbates the DDR. **A.** Sensitivity of *S. cerevisiae* WT, *steΔ* and *tagΔ* cells to the indicated genotoxic agents as assayed by 10-fold serial dilutions of exponentially growing cultures spotted onto YPD plates supplemented with the indicated agents. Plates were incubated at 30 °C from 2 to 5 days. **B.** Exponentially growing *S. cerevisiae* WT, *steΔ* and *tagΔ* cells were treated with 100 µg/mL zeocin and samples collected at the indicated timepoints for Western blot analysis and cytometry (Figure S1D). The activation of the DDR was monitored following the progressive phosphorylation of its downstream effector kinase Rad53. The unphosphorylated and phosphorylated isoforms of Rad53 are indicated (Rad53 and Rad53-P, respectively). Ponceau staining is shown as a loading control. The percentage of Rad53-P was quantified at each point by dividing the raw signal of the upper band by the total signal in that lane, and is plotted in the graph shown on the right. The plotted values represent the mean value of at least 3 independent experiments and the variation is represented by the standard error of the mean (SEM). Unpaired t-tests were used to compare the potential differences of the means at each time point. Only the *p*-value(s) for those being significantly different are indicated. **C.** Identical to (B) but comparing unloaded WT and WT pre-loaded for 2 hours with 0.05% oleate in order to inhibit sterol esterification. **D.** Non-confluent RPE-1 and Huh-7 human cell lines were either left untreated, or treated during 2 hours with the DNA-damaging agents MMS (0.005%) or zeocin (10 µg/mL). Prior to that, cells were pre-loaded with oleate (4 hours at 60 µM) in order to inhibit sterol esterification or treated with 30 µM BSA (mock) as control. The downstream effector kinase of the DDR CHK2 was monitored by Western blot for its activation using a specific antibody against its phosphorylation at Thr68 (P-CHK2). The Western blot signal for total CHK2, calnexin and the Ponceau staining are used as loading controls. One illustrative experiment out of at least 3 per cell type is shown. The mean value of the P-CHK2 to CHK2 signals ratio for genotoxins-treated cells is indicated under each lane. **E.** Identical to (D) though only in response to zeocin and by inhibiting sterols with the specific inhibitor avasimibe (5 µM for 2 hours). The P-CHK2 signal fold-change comparing zeocin + avasimibe *versus* zeocin alone is plotted at the bottom and compiles the values for all experiments for both cell lines.

To gain understanding about the role of STEs in DSB tolerance, we first monitored the progressive phosphorylation of the DDR effector kinase Rad53 upon zeocin treatment as a readout of the DDR activation. *steΔ* cells displayed a modestly faster phosphorylation of Rad53 compared to *tagΔ* and WT cells (Figure 2B), which was unrelated to changes in cell cycle distribution (Figure S1D, lower panel). This phenotype was specific to the DSB-inducing agent zeocin, as HU treatment did not accelerate the pattern of phosphorylation of Rad53 in *steΔ* cells (Figure S1F). To independently validate that this was related to the inability of the cells to esterify sterols, we repeated the experiment in cells pre-loaded with oleate, because this unsaturated fatty acid is firmly characterized to directly inhibit sterol esterification in *S. cerevisiae* ^27^ and outcompetes esterified sterol storage in cultured human cells ^28, 29^. Addition of oleate readily filled LD by increasing TAG production (Figure S1F, upper panel). Cells pre-loaded with oleate also showed a faster Rad53 phosphorylation compared to mock-treated cells (Figure 2C) in the absence of any apparent cell cycle alteration that could explain this difference (Figure S1F, lower panel).

We also explored whether this phenomenon was observable in two different human cell lines, RPE-1 (hTERT-immortalized cells from Retinal Pigmented Epithelium of non-tumoral origin) and Huh-7 (derived from hepatocarcinoma). We induced DSBs with zeocin and the DNA alkylating agent methyl methanesulfonate (MMS) (Figure S2A, upper panels). As a readout for the DDR activation, we monitored the phosphorylation on Thr68 of the downstream effector CHK2. As expected, incubation of RPE-1 and Huh-7 cells with zeocin or MMS led to the phosphorylation of CHK2, while total CHK2 levels remained constant (Figure 2D). Pre-treatment of both RPE-1 and Huh-7 cells with oleate to prevent sterol esterification within LD led to an increased amount of TAG-containing LD (Figure S2B). We observed a stronger P-CHK2 signal in oleate pre-loaded cells compared to mock-treated cells in response to both zeocin and MMS (Figure 2D), suggesting a stronger DDR activation. Moreover, specific inhibition of sterol esterification using the drug avasimibe (Figure S2C) also led to an increased DDR (Figure 2E). Importantly, these phenomena were again unlikely related to cells being in different cell cycle stages, since the short duration of the treatments did not trigger major differences in cell cycle distribution (Figure S2D). Last, in Huh-7 cells, in which zeocin delayed proliferation without abolishing it, preincubation with oleate further sensitized them to zeocin (Figure S2E, right panel).

Altogether, our results show that the inability to esterify sterols and subsequently store STEs in LD leads to an accelerated or exacerbated activation of the response to DSBs in both *S. cerevisiae* and human cells, which consequently impacts cell proliferation.

### Lack of sterol esterification inhibits long-range resection, compromising downstream DSB repair

The esterification of sterols is the consequence of removing free sterols from their previous location, the membrane. To evaluate whether removal of free sterols from membranes was important to control the activation kinetics of the DDR, we used a mutant strain of *S. cerevisiae* deficient for the protein Yeh2 ^30^. Yeh2 is a steryl ester lipase located at the plasma membrane. Its active site being oriented towards the extracellular space, thus with no access to the intra-cellular pool of esterified sterols, it is purported to replenish the free sterol pool of the plasma membrane by hydrolyzing extracellular steryl esters ^31^. In this context, the incapability for maintaining the pool of free sterols at the plasma membrane forces intracellular pools of steryl esters to be hydrolyzed, and the released steryl moieties transported from the lipid droplets and the endoplasmic reticulum to the plasma membrane^32^. As this phenocopies the inability to esterify and store sterols within LD of the *steΔ* strain, we monitored the activation kinetics of the DDR in response to zeocin in isogenic WT and *yeh2Δ* cells and found that, similar to lack of sterol esterification (Figure 2B,C), Rad53 became phosphorylated more rapidly in the absence of Yeh2 (Figure 3A and ^33^) in a cell cycle-unrelated manner (Figure S3A). Again, this was related to DSBs, since the kinetics of Rad53 phosphorylation in response to HU was not accelerated in *yeh2Δ* cells with respect to the WT (Figure S3B). Thus, sterol removal from membranes and their subsequent esterification and storage within LD tone down the speed and intensity of the DDR activation upon DSBs.

**Figure 3.**
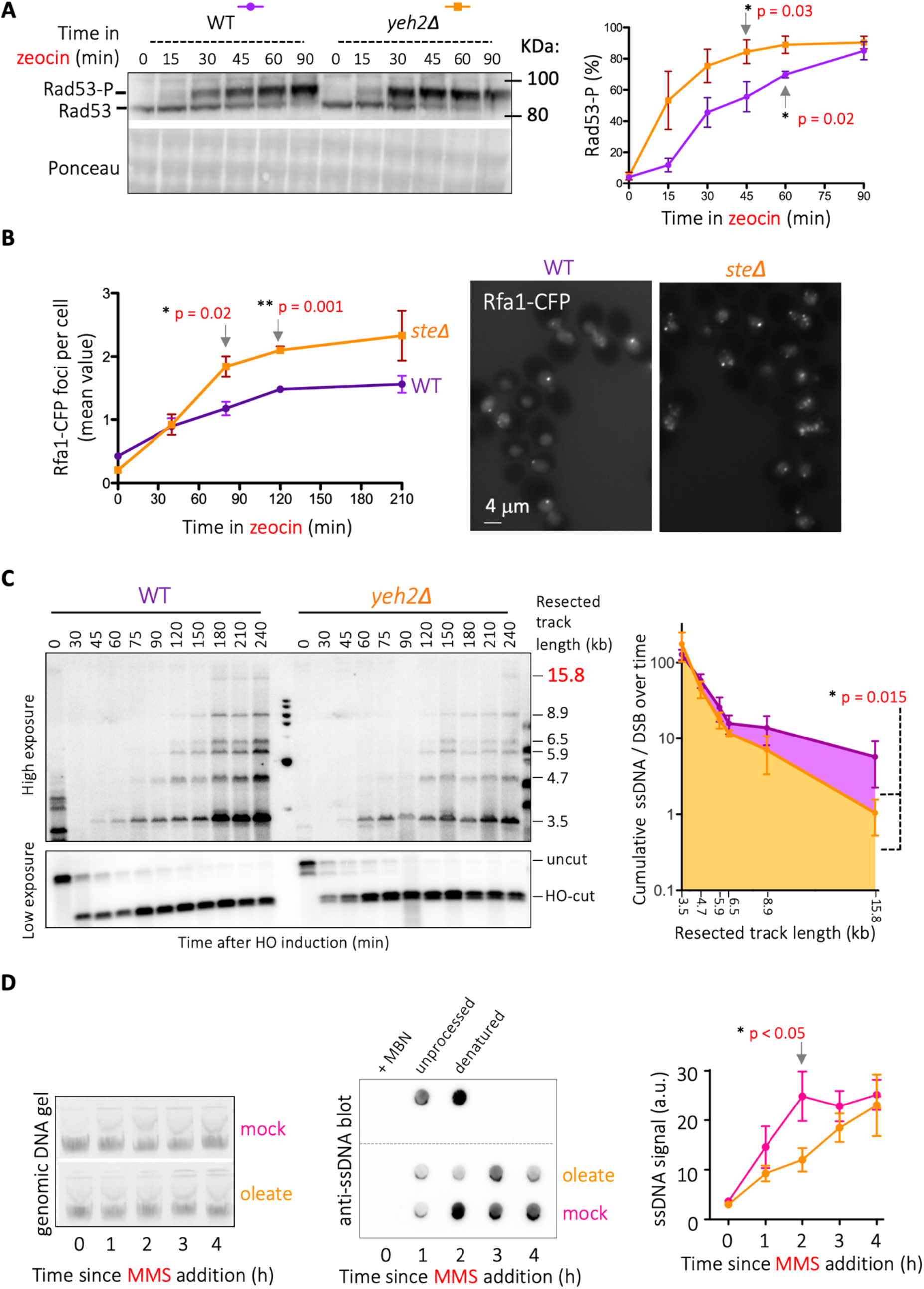
Impairment in sterol esterification blocks processive (long-range) resection. **A.** Exponentially growing *S. cerevisiae* WT and *yeh2Δ* cells were treated with 100 µg/mL zeocin and samples retrieved at the indicated timepoints for Western blot analysis and cytometry (see Figure S3A). The activation of the DDR was monitored following the progressive phosphorylation of its downstream effector kinase Rad53. The unphosphorylated and phosphorylated isoforms of Rad53 are indicated (Rad53 and Rad53-P, respectively). Ponceau staining is shown as a loading control. The percentage of Rad53-P was quantified at each timepoint and plotted in the graph shown on the right. The plotted values represent the mean value of at least 3 independent experiments and the variation is represented as the SEM. Unpaired t-tests were used to compare the potential differences of the means at each time point. Only the *p*-value(s) for those being significantly different are indicated. **B.** Exponentially growing *S. cerevisiae* WT and *steΔ* cells were treated with 100 µg/mL zeocin and samples retrieved at the indicated timepoints for inspection by fluorescence microscopy. The establishment of DNA resection factories was assessed by counting the number of Rfa1-CFP foci per cell. **Left:** the mean values of each timepoint (as calculated in Figure S3D) of three independent experiments were used to obtain this graph (mean and SEM). Unpaired t-tests were used to compare the potential differences of the means at each time point. Only the *p*-value(s) for those being significantly different are indicated. **Right:** representative images of both strains at timepoint 120 min. **C.** WT and *yeh2Δ* cells carrying a plasmid bearing the HO nuclease gene under a galactose-inducible promoter were exposed to 2% galactose to trigger the expression of the HO nuclease to cut at the *MAT locus*, and samples were retrieved at the indicated time points for genomic DNA extraction. Subsequent *Ssp*I digestion, in combination with the use of a probe targeted to *MAT*, allows defining the fate of the cut fragment by Southern blot analysis. As resection progresses on DNA, *Ssp*I restriction sites are lost, leading to progressively longer ssDNA fragments (whose sizes are indicated at the right of the gel) that can be separated on an agarose gel under denaturing conditions. Given the much stronger signal of the “cut” fragments with respect to the “resected” fragments, the gel has been split into low and high contrast halves, respectively. The ssDNA over cut yet unresected DSB molecules was calculated for each shown resection intermediate. The graph on the right plots the sum of all these values (cumulative ssDNA/DSB) for a given resected intermediate during the interval from 90 to 240 min. The error bar is the SEM of three independent experiments. * derives from applying a *t*-test that compares the two populations of values. **D.** Cells were pre-loaded with 60 µM oleate 30 µM BSA (mock) for 4 h, and then exposed to 0.005% MMS for the indicated times. Total genomic DNA was extracted (left panel) and 1500 ng of each condition were spotted onto a nylon membrane directly. The DNA, after being crosslinked to the membrane, was subjected to detection using an anti-ssDNA antibody (middle panel). The raw ssDNA signals obtained this way were quantified for three independent kinetics. The graph (right panel) shows the mean raw ssDNA signal value and the error bars are the SEM. Unpaired t-tests were used to compare the potential differences of the means at each time point. Only the *p*-value(s) for those being significantly different are indicated. As a control for the specificity of the anti-ssDNA antibody, we used the sample “MMS 3 h” for treatment with Mung Bean Nuclease (MBN), in order to digest single stranded DNA (ssDNA), or for denaturation, by addition of NaOH to a final 0.4 N concentration. These three samples are shown in the top part of the membrane.

We next hypothesized that this exacerbated DDR was indicative of alterations somewhere in the cascade ranging from DSB detection to downstream DNA repair. We set to study different steps from the signaling to the repair events occurring in response to a DSB (Figure S3C and ^34^). We first monitored the resection of DSB ends, an early event of DSB processing required for repair by HR. DSB end resection consists on the degradation of the 5’ DNA strand, generating 3’ single-stranded DNA (ssDNA) tails. These structures are covered by the ssDNA-binding heterotrimeric complex RPA, conformed by Rfa1-Rfa2-Rfa3 in *S. cerevisiae*, which forms nuclear foci ^35–37^.WT cells displayed less than one CFP-tagged Rfa1 focus per cell (0.42, mean value) basally. Foci number per cell progressively increased during exposure to zeocin, having tripled at 210 minutes after treatment onset (1.56, Figure 3B & S3D). In contrast, zeocin treatment increased by more than 10-fold the mean number of Rfa1 foci in *steΔ* cells (2.33 *versus* 0.2, Figure 3B and Figure S3D). These results suggest that cells unable to remove sterols from membranes either accumulate longer ssDNA tracts or, on the contrary, accumulate unproductive events because they are unable to implement processive resection. To discern between these two possibilities, we monitored DSB end resection at the molecular level using a genetically engineered strain in which one single DSB can be induced by the controlled expression of the HO endonuclease ^38^. We observed DSBs after 30 minutes of HO induction. Resection products, detected by the progressive disappearance of restriction sites close to DSB ends (Figure S3E), started to accumulate shortly after (Figure 3C). Importantly, *yeh2Δ* cells displayed a decrease in long resection tracks (Figure 3C, 15.8 kb product). We also treated human RPE-1 cells with MMS and physically monitored ssDNA by extracting genomic DNA, spotting it on a membrane, and blotting it with an anti-ssDNA antibody. This way, we detected a progressive accumulation of ssDNA upon MMS addition (Figure 3D). Pre-loading cells with oleate to prevent sterol esterification led to less ssDNA accumulation in response to MMS (Figure 3D). Overall, data point towards a defect in implementing long-range resection in response to DSBs if sterols cannot be removed from membranes or esterified.

If long-range resection is compromised when sterols remain within membranes, thus the efficiency of downstream DNA repair by HR is expected to decrease. We assessed this possibility using a genetic system in which the number of HR repair products, mostly single strand annealing reactions (dependent on Rad52 upon DSB formation), can be measured. The system, integrated in the genome of *S. cerevisiae*, consists of two directly repeated sequences of a *leu2-k* allele that flank one *ADE2* and one *URA3* copy. Whenever a break occurs in between the two *leu2-k* sequences, the recombination process between the two direct repeats will end up with the loss of the intermediate sequences. Cells having lost the intervening *URA3* marker in such a process can be selected in plates containing 5-Fluoroorotic Acid (5-FOA, for the presence of the enzyme encoded by *URA3* will convert 5-FOA to the toxic compound 5-fluorouracil), and thus the frequency of recombinants calculated ^39^. When WT cells were exposed to zeocin, the induction of DSBs led to an expected increase in the frequency of recombination of up to 15-fold (Figure 4A). Importantly, preventing the esterification of sterols either genetically (*steΔ*) or pharmacologically (with oleate) limited the formation of recombination products up to 3-fold (Figure 4A). Likewise, recombination was induced 7-fold in the DSB-prone mutant *rad3-102* ^40^, but the additional *steΔ* mutation restricted this increase to less than 2-fold (Figure S4A). Thus, the decrease in the frequency of HR products suggests that impairment in sterol esterification is hampering DSB repair. To validate this hypothesis in yet another system, we monitored the accumulation of zeocin-induced DSB in RPE-1 and Huh-7 cells using pulsed-field gel electrophoresis (PFGE). In both cell types, zeocin led to a prominent and similar accumulation of in-gel (i.e. broken) molecules (Figure 4B,C, total in-gel molecules and bottom quantifications). Yet, when sterol esterification was inhibited by oleate, the intensity of the shorter fragments increased (Figure 4B,C, lateral orange *vs* pink curves). This result confirms that, while the same number of lesions is induced, repair is less efficient in oleate-treated cells.

**Figure 4.**
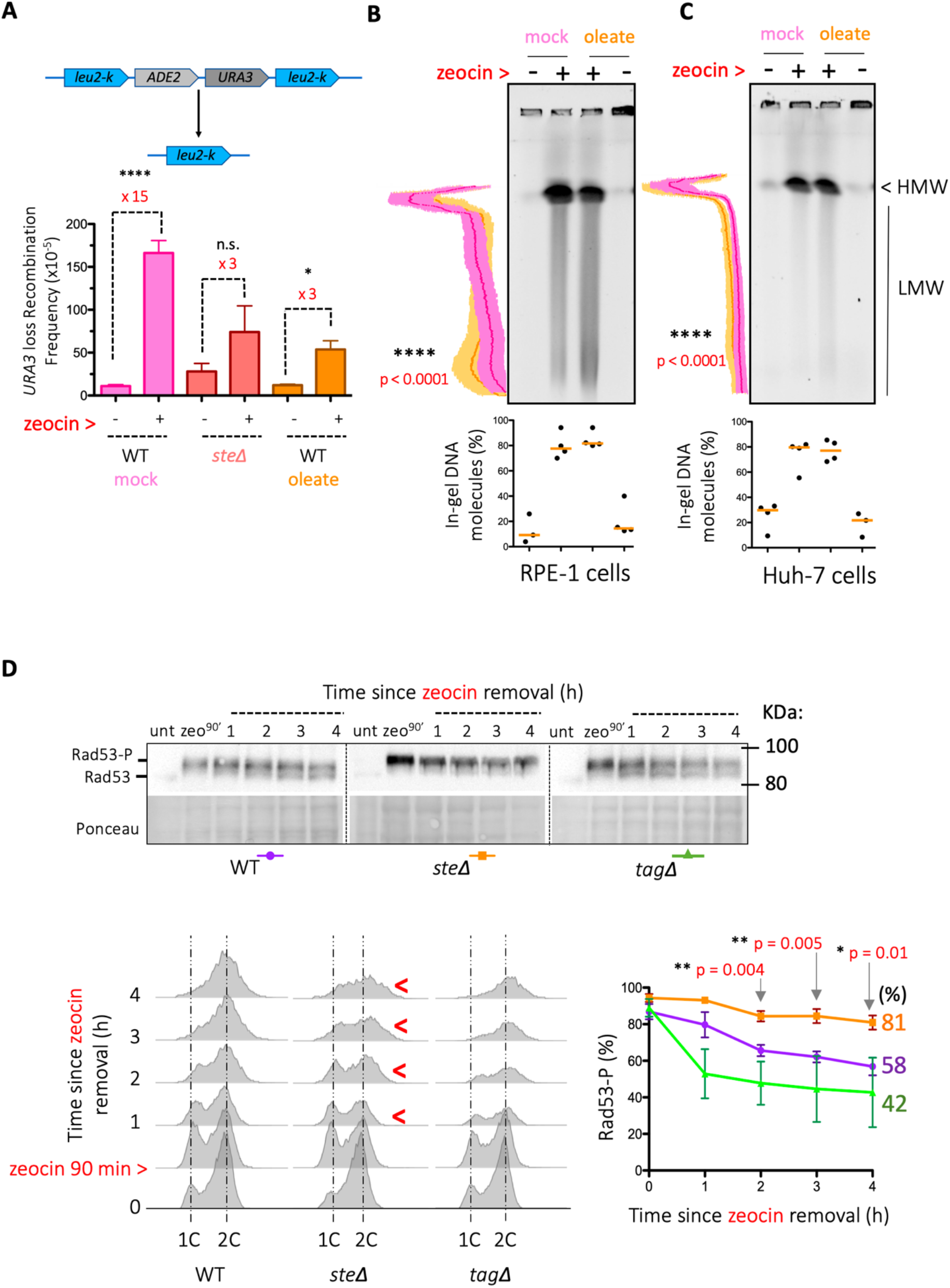
Impairment in sterol esterification prevents DSB repair and DDR de-activation. **A. Top:** scheme of the genome-integrated system allowing the analysis of homologous recombination frequency, mostly through single-strand annealing. In brief, two direct repeats of a *leu2-k* allele flank one *ADE2* and one *URA3* copy. Whenever a break occurs in between the two *leu2-k* sequences, concomitant recombination will end up with the loss of the intervening sequences. This loss of the *URA3* marker can be selected for in plates containing 5-FOA, and the frequency of recombinants calculated. **Bottom:** for each recombination test, WT and *steΔ* cells (for which colonies arousal in +zeocin plates took longer given their sensitivity) streaked onto plates containing 7.5 µg/mL zeocin or not, or 0.05% oleate or not, as indicated, were grown and 6 isolated colonies out of each plate used to measure the number of recombinant and total cells, thus yielding the shown recombination frequencies. For every 6 frequencies derived from 6 colonies, one median frequency would be calculated (= 1 experiment). Each bar represents the mean frequency of the median of at least 3 independent experiments. Zeocin-induced recombination is indicated as a fold-change. Statistical evaluation of the differences between the recombination frequencies was evaluated using a *t*-test: n.s.: non-significant; *: *p*-value < 0.01; ****: *p*-value < 0.0001. **B, C.** Pulsed Field Gel Electrophoresis (PFGE) performed on DNA samples prepared from RPE-1 or Huh-7 cells that had been treated (or not) with zeocin and/or oleate (2 hours at 10 µg/mL zeocin; pre-treatments for 4 hours with 60 µM oleate) in order to evaluate the presence of DSBs under such treatments. Intact DNA molecules remain in the well, while broken molecules migrate into the gel. Given the action of zeocin, in-gel molecules can be subdivided in high-molecular weight fragments (indicated by “HMW”) and low-molecular weight ones (LMW). Plugs were prepared at 37°C to prevent artefactual breaking of DNA molecules ^77^. The bottom graphs provide the percentage of total broken molecules per lane with respect to all molecules in that lane. 3-4 independent experiments are plotted and their median value indicated by an orange bar. The lateral graphs provide the relative intensity (arbitrary units) of signals associated with in-gel broken molecules. The values corresponding to zeocin-treated samples are displayed in pink, those corresponding to oleate + zeocin-treated samples in orange. For pink and orange curves, the mean value of 3-4 independent experiments is displayed in dark colour, while the SEM is indicated in light colours. Statistical analyses of the difference between treatments was assessed using a t-test. **D.** Exponentially growing *S. cerevisiae* WT, *steΔ* and *tagΔ* cells were treated with 100 µg/mL zeocin for 90 minutes. This treatment duration recurrently results in more cells accumulating in the G_1_ phase of the cell cycle (Figure 1B, S1C-D, S3A, 4D, S4B). After 90 min, cells were washed to remove zeocin and resuspended in fresh, zeocin-free medium, and samples were retrieved every hour, as indicated. All samples were processed for Western blot analysis and cytometry. The de-activation of the DDR was monitored following the progressive de-phosphorylation of Rad53. The unphosphorylated and phosphorylated isoforms of Rad53 are indicated (Rad53 and Rad53-P, respectively). Ponceau staining is shown as a loading control. The percentage of Rad53-P was quantified at each time point as in Figure 2B to build the graph shown on the bottom right. The plotted values represent the mean value of at least 3 independent experiments and the variability is represented by the SEM. Unpaired t-tests were used to compare the potential differences of the means at each time point. Only the *p*-value(s) for those being significantly different are indicated. Red arrowheads indicate cytometry timepoints where cell cycle progression was delayed in comparison with the WT strain.

Last, we reasoned that lack of DNA repair would prevent DDR signal extinction even if DSBs are not continuously produced. To test this prediction, we activated the DDR by exposing the cells to zeocin for 90 minutes, removed it from the medium and monitored the de-activation of the DDR over time. WT and *tagΔ* cells showed progressive de-phosphorylation of Rad53 and resumed cell cycle progression (Figure 4D). On the contrary, *steΔ* and *yeh2Δ* cells, respectively unable to esterify sterols or to remove them from membranes, were incapable of Rad53 de-phosphorylation (Figure 4D and Figure S4B) and displayed a delay or a complete halt in cell cycle progression (Figure 4D and Figure S4B, see red arrowheads).

Altogether, the inability to process sterols in response to DSBs leads to an exacerbated DDR, prevents DNA repair and the subsequent de-activation of such a DDR, probably out of a defect in transitioning from short-to long-range resection.

### Lack of sterol processing in response to DSBs concurs with an exacerbated activity of the DDR kinase Tel1/ATM

Tel1/ATM acts at the very first steps of DSB recognition and “short-range” DSB end resection, upon which Mec1/ATR takes over Tel1 in the DDR to activate Rad53 (Figure S3C). But Tel1 participates in the reaction later on by signalling a transient arrest that prevents further “long range” resection (Figure S3C). After this step, Tel1 is displaced by an unknown mechanism, thus endowing DNA repair and cell cycle resumption ^41–43^. Given that we observed both an acceleration in the detection of breaks (Figure 2) and a defect in the transition from short to long range resection (Figure 3), we evaluated if sterol esterification could play a role in the regulation of Tel1 at DSBs. For this purpose, we used a recently created, functional GFP-tagged Tel1 ^44^ to monitor its accumulation in nuclear foci as a readout for its activity at DSBs. In about 40% of the WT cells, 1 to 5 Tel1 foci per nucleus formed after 40 minutes of DSB induction with zeocin, a value that declined down to 20% at 210 minutes (Figure 5A). In contrast, the percentage of *yeh2Δ* cells forming GFP-Tel1 foci doubled that of the WT strain, and *steΔ* cells displaying Tel1 foci reached 70% at the final time point (Figure 5A, right), thus suggesting a higher activity of Tel1 in these mutants. We evaluated whether the level of ATM activation related to the metabolism of sterols in human cells as well. We performed immunofluorescence of ATM phosphorylated at Serine 1981, a marker of its commitment to DNA damage signaling ^45^. When exposing both RPE-1 and Huh-7 cells to MMS, the inhibition of sterol esterification with oleate increased the number of P-ATM-positive cells relative to cells treated with MMS alone, albeit this was not statistically significant (Figure 5B,C, MMS). However, inducing DSBs with zeocin led to a significant increase in P-ATM-positive cells upon inhibition of sterol esterification with oleate (Figure 5B,C, zeocin). Importantly, the increase in the phosphorylation of CHK2 observed when pre-loading the cells with oleate (Figures 2D & 5D) was dependent on ATM activity, as it could be abolished by addition of the ATM inhibitor AZD0156 in both cell types (Figure 5D). Thus, preventing the storage of sterols into LD in response to DSBs leads to an exacerbated activity of the DDR upstream kinase Tel1/ATM.

**Figure 5.**
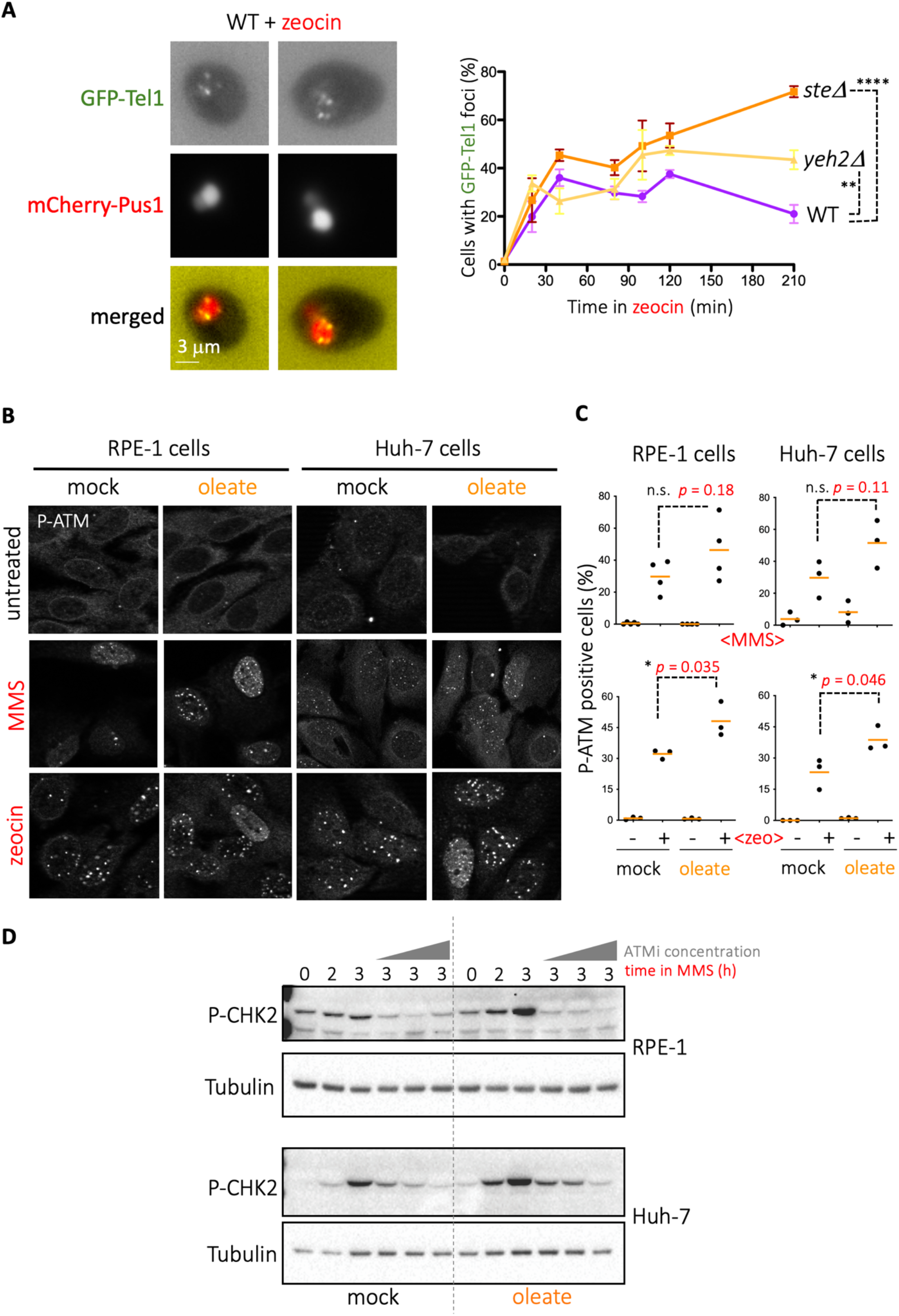
Tel1/ATM displays excessive activation in sterol removal-deficient contexts. **A.** Exponentially growing *S. cerevisiae* WT, *steΔ* and *yeh2Δ* cells were treated with 100 µg/mL zeocin and samples retrieved at the indicated timepoints for inspection by fluorescence microscopy. The involvement of Tel1 in the response to zeocin was evaluated by its ability to form GFP-Tel1 foci. **Left:** illustrative images of the WT strain forming GFP-Tel1 foci in response to 100 µg/mL zeocin after 80 minutes. mCherry-Pus1 was used to define the nucleus boundaries. **Right:** graph displaying the mean value of the percentage of cells forming GFP-Tel1 foci and the SEM of 3 independent experiments for the indicated strains. A one-way ANOVA for multiple comparisons was applied to evaluate whether there were significant differences between the means of the 210 time-point and the obtained *p*-values are indicated by asterisks. **, *p* < 0.01; ****, *p* < 0.0001. **B.** Representative images obtained after immunofluorescence to detect P-ATM (ATM phosphorylation at Serine 1981) signal in RPE-1 and Huh-7 fixed cells that had been treated (or not) with 0.005% MMS or 10 µg/mL zeocin for 2 hours; pre-treatments were done for 4 hours with 60 µM oleate. **C.** Quantification of the percentage of P-ATM-positive cells. The values obtained from 3 to 4 independent experiments are plotted as black spots, and the mean of them is shown as an orange bar. At least 200 cells were counted per condition and experiment. The *p*-values obtained after performing an unpaired *t*-test are shown. **D.** RPE-1 and Huh-7 cells were treated or not with MMS 0.005% for 2 hours. Thereafter, the ATM inhibitor AZD0156 was added at a final concentration of 1, 10 or 50 nM for 1 hour. Cells were collected and phosphorylation on Thr68 of CHK2 and tubulin levels were analyzed by Western blot.

### Tel1/ATM binds Phosphatidylinositol-4-phosphate

Sterols modulate proteins through their transmembrane segments, yet Tel1 and ATM are soluble proteins. We reasoned that sterol presence in membranes could indirectly affect other lipids with regulatory potential, in turn impacting Tel1/ATM activity. Tel1 and ATM are phosphatidylinositol-3-kinases-like kinases (PI3KKs), thus ancestrally related to phosphatidylinositol-3-kinases ^22^. We asked whether Tel1 kept the potential of a physical interaction with phosphoinositides (PIPs). We immunopurified a functional FLAG-tagged Tel1 (Figure S5A,B) from *S. cerevisiae* ^46^ and incubated the eluate with a commercially available membrane on which Phosphatidylinositol (PI) and its six phosphorylated PIP variants are spotted. Additional, unrelated lipid species are also spotted on that membrane. We observed robust anti-FLAG signals specifically against the three spotted monophosphate phosphoinositides, namely PI(3)P, PI(4)P and PI(5)P (Figure S5C), while FLAG-Tel1 affinity for the di-and tri-PIP species was low or absent. In our particular interest, PI(4)P is related to the metabolism of sterols at the interface between the Trans-Golgi and the endoplasmic reticulum (ER). There, the OSBP1 exchanger promotes the extraction of sterol moieties from the ER and their insertion in the Trans-Golgi in exchange of PI(4)P, which then follows a reciprocal fate (Figure 6A) ^15, 47, 48^. Thus, if Tel1/ATM were to bind PI(4)P, this could explain why high level of unesterified sterols affects this kinase. We incubated again purified FLAG-Tel1 with a membrane in which only the three variants of PI(4)P were spotted, and observed again a clear preference for the monophosphate species, while no signal at all was obtained if a similar membrane was incubated with 3XFLAG peptides only (Figure 6B). In both human RPE-1 and Huh-7 cells, we assayed the interaction of ATM with PI(4)P by proximity ligation assay (PLA) using a well-documented anti-PI(4)P antibody ^49^ and an anti-total ATM antibody. The PLA assay yielded specific signals in the cytoplasm (Figure 6C), as demonstrated by loss of signals in technical controls (absence of one primary antibody at a time) (Figure 6C, bottom left), or biological controls (siRNA-mediated depletion of ATM) (Figure 6C, bottom right and Figure S5D). Yet, basal levels detected in Huh-7 cells were low, close to the one-antibody-only control levels, probably indicating a less frequent basal association in these cells (Figure 6C). Thus, Tel1/ATM interacts with PI(4)P in the absence of DNA damage.

**Figure 6.**
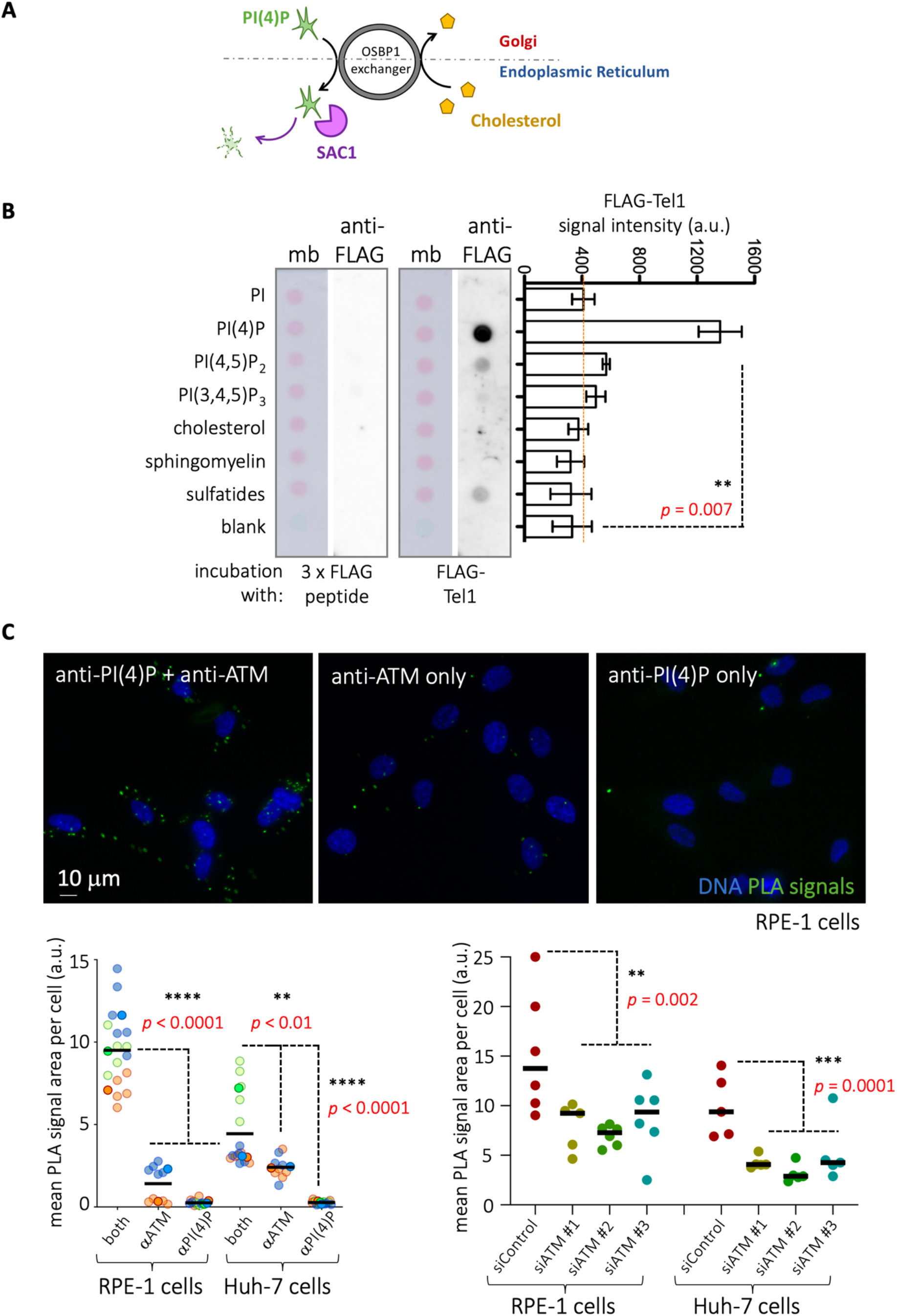
ATM/Tel1 displays a physical link with PI(4)P. **A.** Illustration of how the OSBP1 exchanger working mechanism. The distance between the Golgi and the Endoplasmic Reticulum (ER) is represented by the dashed line and their connection established through OSBP1, whose activity is represented as a wheel whose directionality is given by the black arrows. It extracts cholesterol from the ER and sends it to the Golgi in exchange for PI(4)P that is extracted from the Golgi and sent to the ER. In the ER, the phosphatase SAC1 hydrolyses PI(4)P, which releases ATP, further fostering the exchange. **B.** A hydrophobic membrane on which 100 pmol of the indicated lipid species were spotted was incubated with immunopurified FLAG-Tel1 and further developed using an anti-FLAG antibody. The bar lengths in the graph represent the mean value of 4 independent experiments and the error bars the associated SEM. To assess whether the mean values differed significantly, a *t*-test was performed and *p*-values are shown. Additionally, an identical lipid-containing membrane was incubated with 3xFLAG peptide and subsequently developed using an anti-FLAG antibody to control that the observed signals were not due to FLAG peptide binding alone. PI; phosphatidylinositol; PI(4)P, phosphatidylinositol-4-phosphate; PI(4,5)P, phosphatidylinositol-4,5-bisphosphate; PI(3,4,5)P, phosphatidylinositol-3,4,5-triphosphate. **C.** Proximity ligation assay (PLA) to assess whether ATM and PI(4)P are in close proximity (< 40 nm). **Top:** representative pictures of RPE-1 cells, in which PLA signals are pictured in green. **Bottom left:** the graph shows the mean PLA signal area occupied per cell when doing the experiment with both antibodies (experiment) or by omitting one primary antibody at a time (technical controls). In practice, each point is the value of having measured all the PLA signals present in one photo (20 to 40 cells) and divided this value by the number of nuclei. To account for reproducibility, we used SuperPlots to draw the graphs ^78^: each independent experiment is plotted in a different colour, where the mean value of each independent experiment is highlighted by a more intense colour than the individual values of that experiment, for which the colour is more translucid. Last, the solid horizontal line marks the mean of the means. A one-way ANOVA for multiple comparisons was applied to evaluate whether there were significant differences between the means. **Bottom right:** the graph shows the mean PLA signal area occupied per cell when doing the experiment either in cells transfected with an siRNA Control (experiment) or with three different siRNAs targeted against ATM (biological controls). The meaning of each plotted point, of the horizontal line and the statistical analysis is as explained above. ATM depletion efficiency can be seen in the Western blots presented in Figure S5D. All the experiments have been performed in both RPE-1 and Huh-7 cells.

### ATM/Tel1 nuclear availability is conditioned by the OSBP1 exchanger

The OSBP1 exchanger activity is driven forward thanks to the activity of the SAC1 phosphatase, which continuously hydrolyzes the translocated PI(4)P moieties that reach the ER (Figure 6A, and ^15, 47, 48^). Considering that *i)* Tel1/ATM binds PI(4)P in its basal state (Figure 6) and that *ii)* high level of sterols in ER membranes, which promotes PI(4)P exchange and hydrolysis ^47, 50^, promotes an exacerbated Tel1/ATM activity (Figure 5), we hypothesized that non translocated PI(4)P present at the Golgi may act as a lock that limits Tel1/ATM availability within the nucleus.

To test this hypothesis, we first reasoned that DNA damage, which causes ATM relocation to the nucleus, would decrease the ATM-PI(4)P proximity signals obtained in PLA experiments. In RPE-1 cells, where basal signals were elevated enough (Figure 6C) as to assess a putative decrease, zeocin or MMS treatments significantly led to a loss of ATM-PI(4)P PLA signals (Figure 7A). Moreover, detection of Golgi-associated PI(4)P molecules by means of a transfected fluorescent biosensor demonstrated a reduction in the area occupied by these signals, suggestive of PI(4)P consumption in response to these agents (Figure S6A). Reciprocally, treating cells with the specific ATM inhibitor AZD0156 increased the number of PLA signals (Figure S6B). Thus, it appeared that the association of ATM with PI(4)P inversely correlated with the need for ATM within the nucleus.

**Figure 7.**
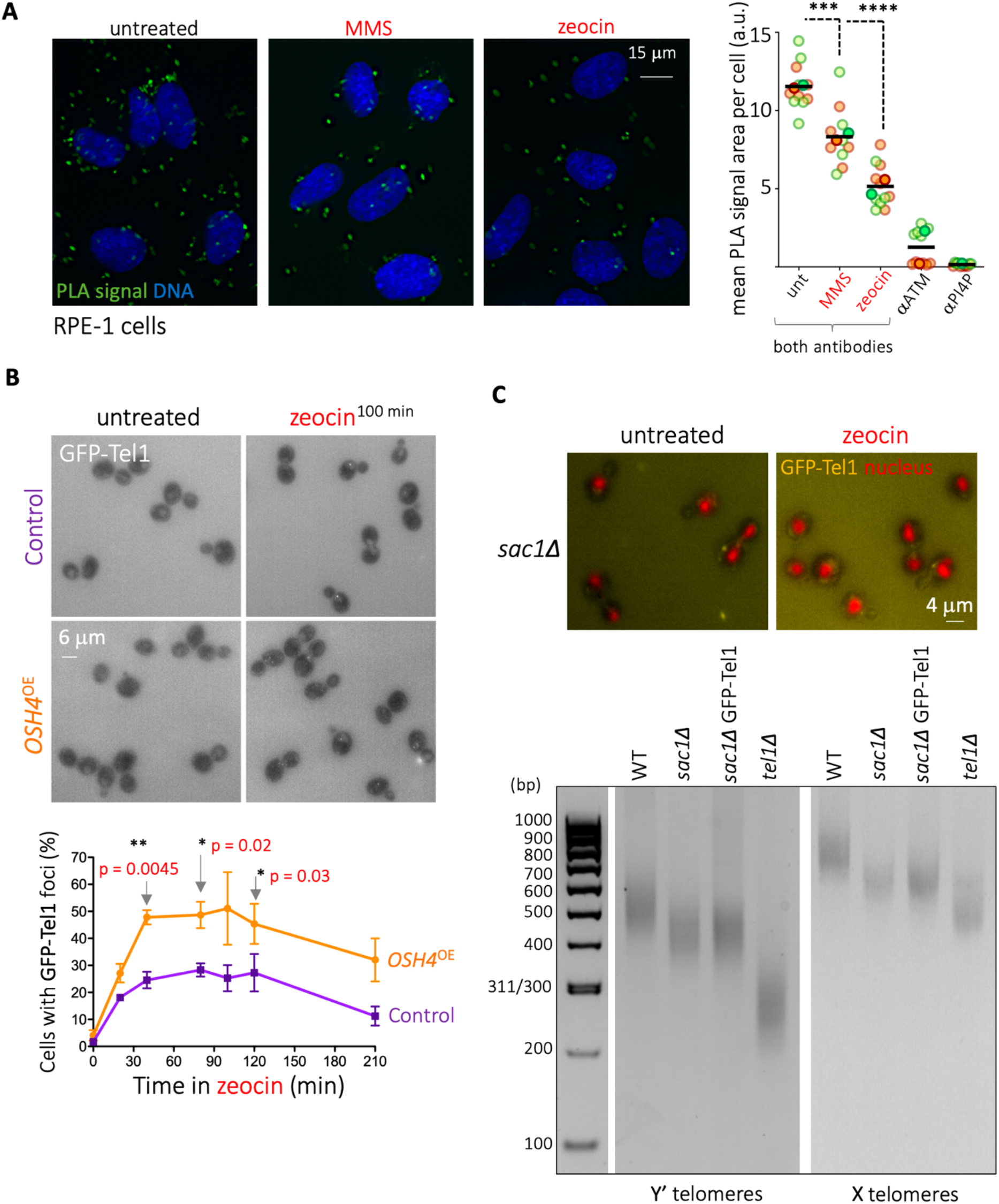
ATM / Tel1 presence in the nucleus is ruled by OSBP1 / Osh4 activity. **A.** PLA to assess whether ATM and PI(4)P proximity is altered in response to DNA damage. **Left:** representative pictures of RPE-1 cells, in which PLA signals are pictured in green, untreated or exposed for 2 hours to MMS (0.005%) or zeocin (10 µg/mL). **Right:** the graph shows the mean PLA signal area occupied per cell, where each point represents the value of all measured PLA signals in one image (20 to 40 cells) divided the number of nuclei. Each independent experiment is plotted in a different colour. The mean value of each independent experiment is highlighted by a more intense colour than the individual values of that experiment, for which the colour is more translucid. The solid horizontal line indicates the mean of the means. A one-way ANOVA for multiple comparisons was applied to evaluate whether there were significant differences between the means of the indicated conditions. **B.** Exponentially growing *S. cerevisiae* WT cells transformed with an empty vector (“Control”) or with a plasmid overexpressing *OSH4* were treated with 100 µg/mL zeocin and samples collected at the indicated timepoints for inspection by fluorescence microscopy to evaluate the percentage of cells displaying GFP-Tel1 foci. **Top:** illustrative images of GFP channel (Tel1 signals) in the indicated conditions. **Bottom:** graph displaying the mean value of the percentage of cells forming GFP-Tel1 foci and the SEM of 3 independent experiments for the indicated conditions. Unpaired t-tests were used to compare the potential differences of the means at each time point. Only the *p*-value(s) for those being significantly different are indicated. **C. Top:** mCherry-Pus1 GFP-Tel1 *sac1Δ* cells were exposed to 100 µg/mL zeocin for 2 hours or not and imaged to establish whether Tel1-associated signals emanated from the nucleus and / or the cytoplasm. **Bottom:** Telomeres (X and Y’) length was measured by PCR-mediated amplification (see Materials & Methods) from genomic DNA extracted from the indicated strains. *tel1Δ* cells were included as a control given their shortened telomere phenotype. Note that GFP-tagging Tel1 does not influence telomeres length.

Second, we surmised that, since boosting the activity of the OSBP1 exchanger consumes PI(4)P more readily, this should render more ATM/Tel1 available for the nucleus. We overexpressed the OSBP1 counterpart in *S. cerevisiae*, Osh4 ^47, 51^, then exposed cells to zeocin and monitored their ability to form nuclear Tel1 foci in comparison with cells expressing an empty plasmid. While the kinetics of Tel1 foci formation remained unchanged, the percentage of cells displaying nuclear Tel1 foci was increased 20 to 30% throughout the whole kinetics when Osh4 was overexpressed (Figure 7B), in agreement with our prediction.

Third, in a reciprocal approach, we reasoned that, if cells could not hydrolyze PI(4)P, such as in the absence of SAC1, PI(4)P is expected to accumulate thus sequestering Tel1 away from the nucleus. In agreement, GFP-Tel1 foci did not form in the nuclei of *sac1Δ* cells, even in response to zeocin treatment, and fluorescent signals would emanate from cytoplasmic structures only (Figure 7C, upper panel, yellow cytoplasmic signals). Additionally, as the absence of Tel1 leads to shortened telomeres ^52^, we used a terminal transferase-assisted PCR ^53, 54^ to assess the length of telomeres in a strain deleted for *SAC1*. Indeed, we observed that both X and Y’ telomeres display an intermediated length in *sac1Δ* compared to WT and *tel1Δ* cells (Figure 7C, gel). Altogether, we conclude that ATM/Tel1 availability in the nucleus is conditioned by its binding to cytoplasmic PI(4)P.

### The OSBP1 exchanger conditions ATM activation and activity

We then wanted to ascertain whether the reduced availability of ATM in the nucleus as shown above translated into functional consequences when cells were exposed to different types of agents creating DSBs. To assess this, we treated cells with different ATM-activating genotoxins then monitored whether phosphorylated ATM signals could be switched off by adding the specific OSBP1 inhibitor schweinfurthin G (SWG) ^55^. As a control, we did the same experiments using the ATM inhibitor AZD0156 instead of SWG. First, we recurrently observed that genotoxic treatments led to the stabilization of ATM protein (Figure S7A). As such, we turned to use calnexin as a loading control (Figure 8A & S7A). Although there were subtle differences when comparing the various genotoxins, inhibiting OSBP1 had the ability to reduce phosphorylated ATM levels in RPE-1 cells in all instances (Figure 8A). Thus, the predicted decrease in ATM availability in the nucleus matched a decreased level of activated ATM.

**Figure 8.**
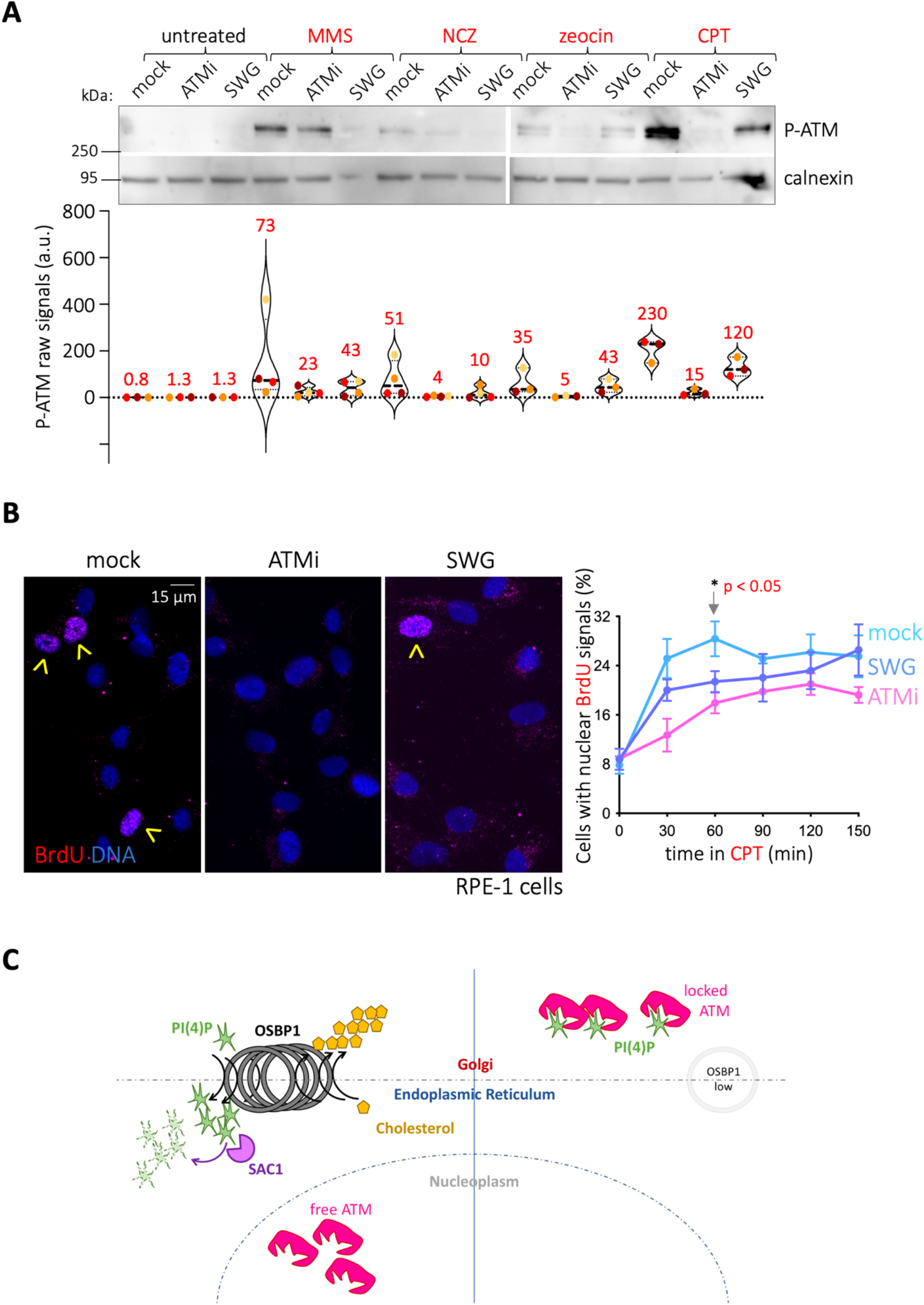
OSBP1 inhibition limits ATM activation. **A.** Western blot showing P-ATM (ATM phosphorylation at Serine 1981) detection. Calnexin is used as an orientative loading control (see Figure S7A for explanation). The experiment assesses the ability of the ATM inhibitor AZD0156 (ATMi) or the OSBP1 inhibitor schweinfurthin G (SWG) to extinguish P-ATM signals triggered by the previous exposure to different genotoxins. Thus, RPE-1 cells were treated either with 0.005% MMS, or 10 µg/mL zeocin, or 120 ng/mL neocarzinostatin (NCZ), or 200 µM camptothecin (CPT, see Materials & Methods) for 2 h, then 500 nM AZD156 (ATMi) or 10 nM SWG were added in the sustained presence of the genotoxins for 2 additional hours. The images are illustrative of three to four independent experiments whose quantifications are shown in the bottom. Each point illustrates the observed raw P-ATM signal for each condition in each experiment, and similarly coloured dots belong to a same experiment. All the signals belonging to a same experiment were obtained from simultaneously run and transferred gels and simultaneously hybridized and exposed membranes. Thick dashed black lines represent the median of the experiments and their specific value is shown as a red number on top of each violin plot. **B.** RPE-1 cells were grown with 10 µM BrdU in the culture medium for the last 16 h prior to further processing. Cells were then either mock-treated, or pre-treated for 1 h with 500 nM of the ATM inhibitor (ATMi) AZD0156 or 10 nM of the specific OSBP1 inhibitor schweinfurthin G (SWG). Then, 200 µM CPT was added and cells were processed for BrdU detection (see Materials & Methods) at the indicated time points. **Left:** illustrative images of BrdU-positive nuclei, indicated by yellow arrowheads. **Right:** the percentage of cells in the population whose nuclei were positive for BrdU was established by visual inspection of the acquired images. The graph shows the mean percentage out of three independent experiments and the error bars indicate their associated SEMs. Unpaired t-tests were used to compare the potential differences of the means at each time point. Only the *p*-value(s) for those being significantly different are indicated. **C. Model:** The constitutive binding of ATM/Tel1 to PI(4)P presumably takes place at the Golgi and limits the molecules of ATM/Tel1 that are available to operate in the nucleus. **Left:** conditions enhancing the activity of the OSBP1/Osh4 exchanger, such as elevated (chole)sterol levels in the ER or increased activity of the phosphatase SAC1/Sac1, promote PI(4)P extraction from the Golgi and its hydrolysis at the ER. This releases ATM/Tel1, which can then reach the nucleus and act in DNA-related transactions, such as DNA DSB signalling and telomere length maintenance. **Right:** low (chole)sterol availability in the ER, lack of PI(4)P hydrolysis in the absence of SAC1/Sac1, or OSBP1 inactivity cause PI(4)P accumulation in the Golgi, which sequesters ATM away from its nuclear functions.

Last, and as a readout for events downstream of ATM signaling, we evaluated whether prior inhibition of OSBP1 with SWG had an impact on the onset of DNA resection, as expected if ATM activation (and activity) are delayed or reduced. We incubated cells with the thymidine analogue BrdU for 16 h. BrdU incorporated in double-stranded DNA can only be detected by an anti-BrdU antibody if resection starts, thus exposing it ^56^. Since the kinetics of BrdU exposure is well established in response to camptothecin (CPT) ^57^, we exploited this framework in our experimental setting. In RPE-1 cells, nuclei displaying resection could readily be detected at 30 min and their number peaked 60 min after CPT addition, remaining stable from that moment on (Figure 8B). In marked contrast, specific ATM inhibition by AZD0156 sharply diminished (yet without abolishing) both the kinetics and the maximum number of nuclei engaged in resection (Figure 8B). OSBP1 inhibition by SWG similarly slowed down the kinetics with which cells engaged in resection (Figure 8B). The effects observed in Huh-7 cells followed the same trend as described although once more in a less robust manner (Figure S7B). Thus, the availability and, concomitantly, the activity of ATM/Tel1 in the nucleus is dependent on the activity of the OSBP1 exchanger.

## Discussion

In this work, we report that the physical and functional availability of ATM/Tel1 in the nucleus is subordinated to its cytoplasmic binding to the lipid PI(4)P, whose half-life is itself linked to the activity of the OSBP1 exchanger. In response to DNA damage, ATM/Tel1 dissociates from PI(4)P. At the same time, the damage elicits the esterification and storage of sterols towards LD, which in turn reduces the level of free sterols in the ER. This establishes a negative feed-back that slows down PI(4)P extraction from the Golgi and its subsequent hydrolysis, eventually contributing to the titration of ATM/Tel1 away from the nucleus. This dampening is critical for the transition from short to long-range resection and downstream DNA repair, as well as may be key in ruling the potential of cells for adaptation. Furthermore, we demonstrate that manipulating the sterol-PI(4)P axis permits an unanticipated control of ATM/Tel1 (Figure 8C, model).

### PI(4)P is a crucial piece in the puzzle of ATM biology

In its inactive state, ATM displays a dimeric conformation and, upon activation, its autophosphorylation at serine 1981 correlates with its monomerization. While this autophosphorylation is essential *in vivo* to activate ATM ^45^, its function remains elusive because it is completely dispensable *in vitro*. In this case, dimer to monomer transition as well as substrate recognition and phosphorylation by ATM still occurs without autophosphorylation ^58^. Previously, it was suggested that a yet undiscovered factor may bind ATM basally, thus preventing its monomerization, and thereby inhibiting its kinase activity ^59^. Herein, we show that ATM/Tel1 associates constitutively with PI(4)P, and that PI(4)P consumption is necessary to promote ATM translocation to the nucleus in response to DNA damage. Furthermore, we show that cell treatments with genotoxic drugs leading to ATM activation elicit the dissociation of ATM and PI(4)P. We therefore propose that PI(4)P is the missing factor that, *in vivo*, keeps ATM inactive. It would be important to asses, in future work, how ATM autophosphorylation at serine 1981 and PI(4)P binding relate to each other.

Our data also shed light on another mystery surrounding Tel1/ATM biology. Tel1 and ATM are both reported to regulate the transition from short- to long-range resection at DNA double strand breaks, but the mechanism controlling Tel1/ATM unbinding from DNA to promote resection resumption is unknown ^41, 42^. Here, we propose that the removal of sterols from ER membranes via their storage within LD, as induced by DNA breaks, slows down the OSBP1 exchanger activity, thus decreasing PI(4)P hydrolysis. PI(4)P levels increased this way progressively sequesters ATM away from the nucleus. The window of action of ATM would therefore depend on the time needed for sterol levels to drop at the ER. This implies that the ER is a key organelle in the attenuation of the DDR, thus ruling crucial downstream aspects such as DNA repair, adaptation to DNA damage or apoptosis. Additionally, our model implies that, in response to DNA damage, the specific esterification and storage of sterols within LD needs to be instructed. It will be key in the future to assess whether the DDR itself targets factors involved in this process.

### Therapeutic implications of the DDR-lipids connection

Cancer cell proliferation is supported by attenuation of the DDR and increased lipid metabolism. Cancer cells bear an increased number of LD that correlates with tumor aggressiveness, bad prognosis, and chemotherapy resistance ^11–13, 60, 61^. Our work identifies the cholesterol-PI(4)P axis as a novel key Tel1/ATM activity controller, able to govern the kinetics and intensity of the DDR in response to DSBs. We propose that cancer cells may maintain their typically irresponsive DDR through a modification of the sterol-PI(4)P-ATM loop warranting them an attenuation of their DDR. This could happen through an enhanced storage of sterol esters within LD. Reminiscently, an increase in the number and size of LD had already been reported in *mrx* mutants, deficient in DSB sensing and repair ^62^. DDR attenuation could also occur, as recently reported, by an exacerbated synthesis of PI(4)P, to which adenocarcinoma cells are addict ^63^. In support, recent findings highlight the potential of targeting the metabolism of cholesterol for modulating the DDR in gallbladder cancer ^64^.

Our findings could also be important for non-dividing cells, such as post-mitotic neurons. In these cells, a low ATM activity drives DSBs repair by non-homologous end joining ^1, 65^, and a precise control of this DDR is key ^66^: an excessive activity of ATM de-compacts chromatin ^67^, and open chromatin becomes more prone to repair by HR ^68^. Consequently, the excessive activity of ATM forces the progression of these cells to S phase, which triggers apoptotic mechanisms in neurons ^2, 69–71^. Thus, limiting the extent of ATM activity by the lipid environment in a non-cycling cell may be crucial for its survival.

Last, our findings could shed light onto mechanisms underlying lipodystrophy, where a loss of white adipose tissue gives rise to ectopic fat deposition and engenders severe metabolic alterations. The lack of adipose tissue can be ascribed either to incorrect adipocyte development, or lack of TAGs formation / storage within LD. Yet, in the case of some familial partial lipodystrophies, such as Hutchinson-Gilford Progeria Syndrome (HGPS), the causative molecular defect has remained poorly linked to problems in LD storage. Molecularly, HGPS resembles a genome instability syndrome: an irreversibly anchored Lamin A destroys the integrity of the nuclear envelope, further leading to altered gene expression, DNA replication, and telomere maintenance, thus strongly activating the DDR and hampering downstream DNA repair ^72^. In view of our work, these features and the lipodystrophic phenotype become suddenly linked: the strong DDR activation would be permanently paralleled by sterol esterification and storage, which would override TAG storage. Consequently, TAG displacement may lead to their redistribution within the body and loss of adipocyte identity. In support of this idea, mutations in genes related to genome integrity preservation, and which also elicit a strong DDR, such as in the Werner helicase, or in the SMC5/6 sumo ligase NSE2, also lead to lipodystrophy ^73, 74^, a fact that has remained puzzling since these genes are completely unrelated to the metabolism of lipids.

### Concluding remarks

We uncover a link between two previously poorly connected research fields. This raises a myriad of issues. For example, it is generally believed that sterol esterification is activated mostly when transport mechanisms are saturated, thus making the level of free sterols in the ER reach a high concentration ^75^. We propose here that this process is actively instructed in response to DSBs, thus raising the need to unveil the DDR effectors in charge of LD formation. It would also be key to assess how and when LD are dismantled in order to respond to new waves of DNA damage.

The overall effects we report are mild given that the metabolism of sterols is finely controlled in a multilayered manner ^75^, thus permitting a limited window of experimental manipulation. Yet, our data unveil the existence of an evolutionarily conserved mechanism from yeast to human cells linking lipid biology and the response to DSBs. Crucially, the pharmacological manipulation of this mechanism allows us to explore new therapeutic interventions. Overall, this work paves the road for abundant, novel and exciting research.

---

**Figure S1.**
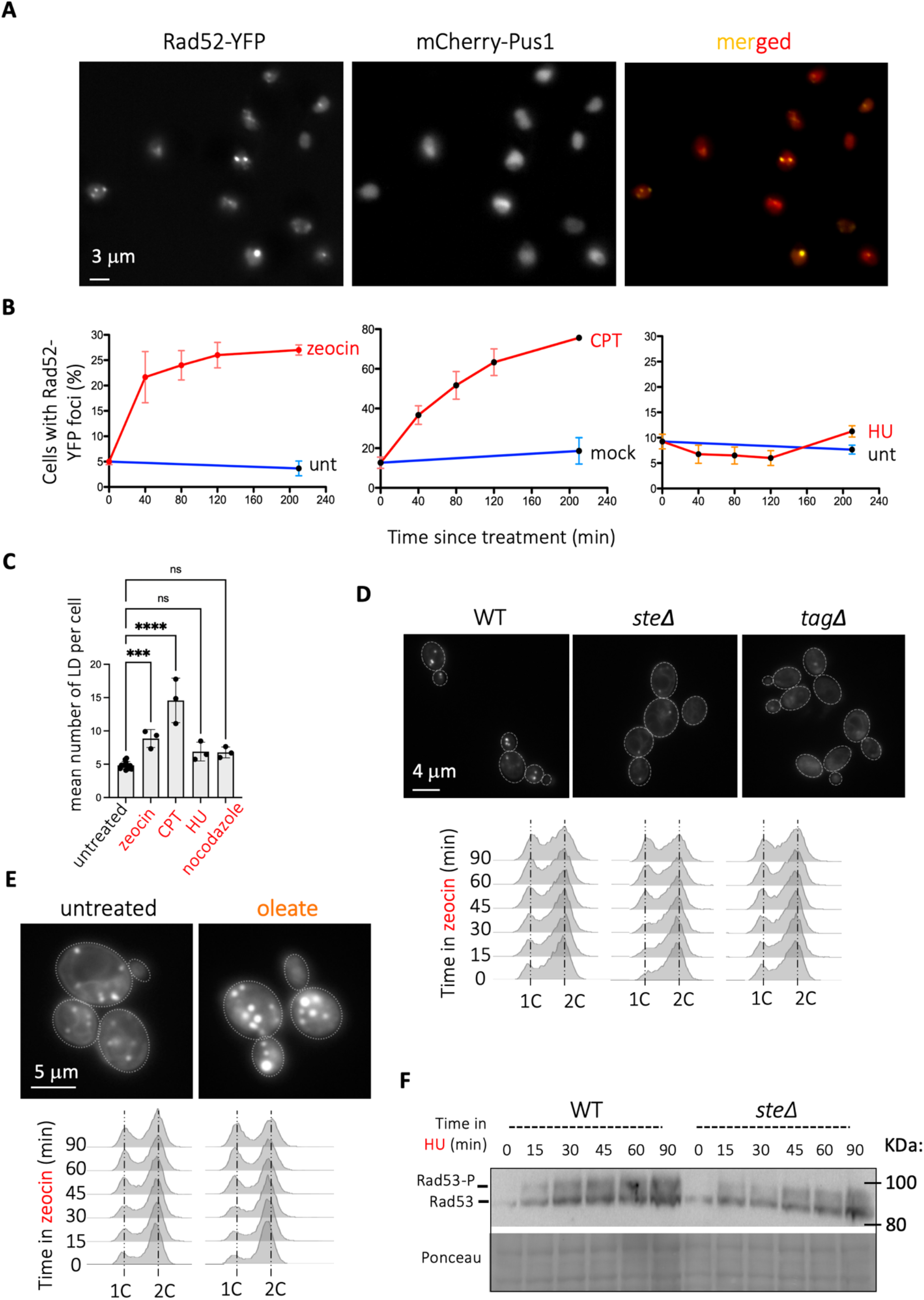
Controls for Figures 1 and 2. **A.** Representative image to illustrate the Rad52 foci (Rad52-YFP panel) in *S. cerevisiae* cells that were exposed to 100 µg/mL zeocin. mCherry-tagged Pus1 was used as a tool to precisely define the nucleus throughout all the experiments presented in Figure S1B. **B.** The percentage of cells bearing Rad52-YFP foci at each time point of the experiments described in Figure 1 was inspected visually. Each plotted value harbours the information referring to at least 3 independent experiments, each with at least 150 cells. The variation is indicated by the standard error of the mean (SEM). Red lines are used for drug-treated cultures, while blue lines are used for untreated ones. **C.** The mean value of LD per cell at time 210 min of treatment (or associated untreated controls) as counted from each of the independent experiments done to build Figure 1B-E were plotted together in order to perform a statistical analysis of the potential difference among the means of the means. To this end, a multiple comparisons one-way ANOVA was applied. ***, *p* < 0.001; ****, *p* < 0.0001; ns, non-significant. **D. Top:** Nile Red staining of cells to verify that the *steΔ* and *tagΔ* strains bear less LD and of decreased intensity, as reported ^76^. **Bottom:** cytometry profiles concerning the experiments presented in Figure 2B. **E. Top:** example of the systematic verification prior to the experiment presented in Figure 2C: Nile Red staining to ensure that the cells had indeed taken the provided oleate from the culture medium and stored it in the shape of TAGs within LD. **Bottom:** cytometry profiles concerning the experiments presented in Figure 2C. **F.** Identical experiment as that reported in Figure 2B but in response to 20 mM hydroxyurea (HU).

**Figure S2.**
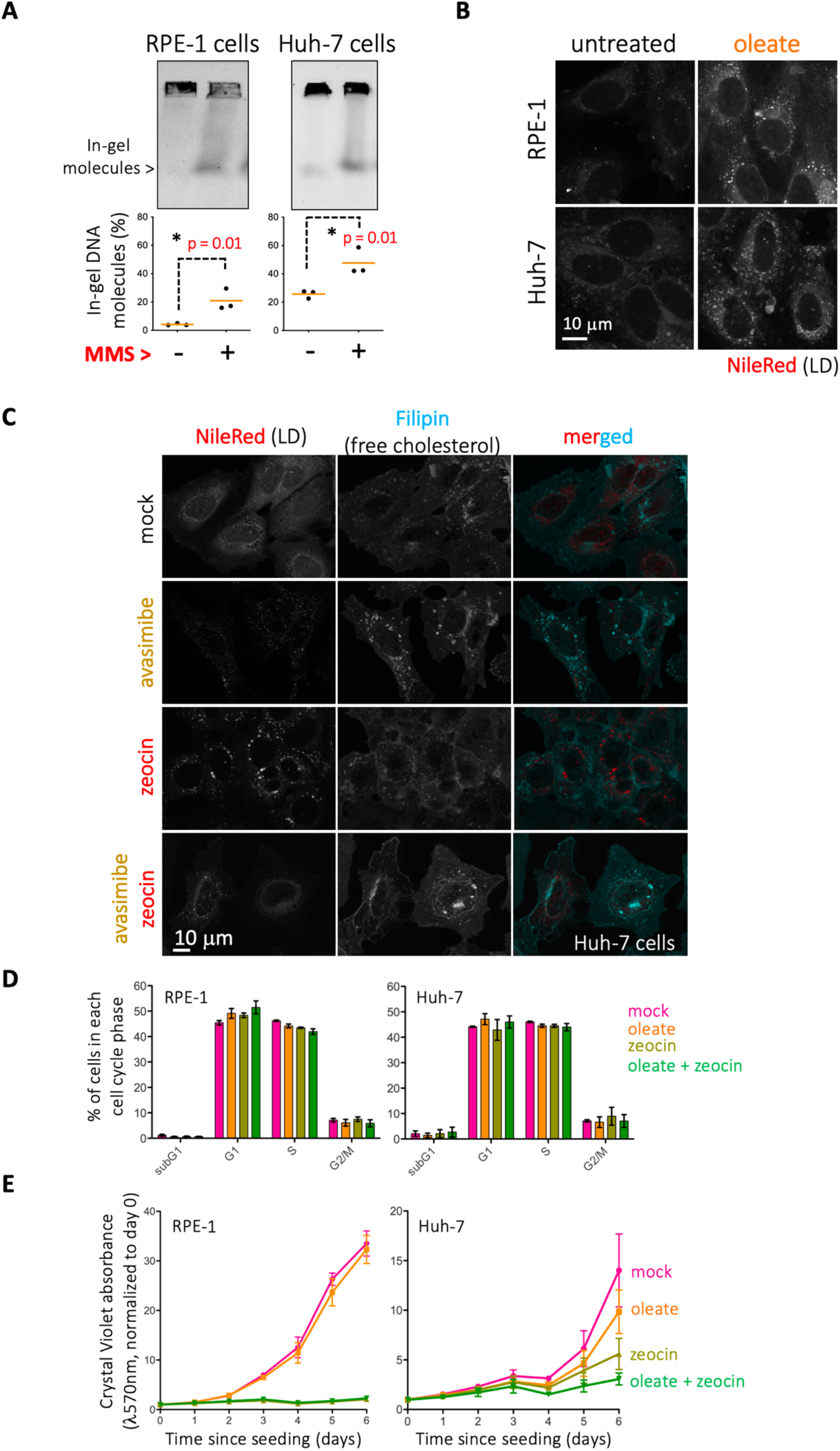
Additional controls for experiments presented in Figure 2. **A.** Pulsed Field Gel Electrophoresis (PFGE) performed on DNA samples prepared from RPE-1 or Huh-7 cells that had been treated (or not) with 0.005% MMS for 2 hours to evaluate whether this agent creates DSBs in these cells. Intact DNA molecules are retained in the well, while broken ones migrate into the gel (indicated as “in-gel molecules”). The bottom graphs provide the percentage of broken molecules (with respect to all molecules in a given lane) per lane. Plugs were prepared at 37°C to prevent artefactual breaking of DNA molecules ^77^. Three independent experiments are plotted and their mean value is indicated by an orange bar. The statistical significance upon a *t*-test is indicated as the *p*-value. **B.** Representative images of fixed Huh-7 and RPE-1 cells dyed with Nile Red during the systematic verification of the experiments presented in Figure 2D, in order to ensure that the cells had incorporated the provided oleate from the culture medium and stored it in the shape of TAGs within LD. **C.** Representative images of fixed Huh-7 cells dyed with Nile Red and filipin during the systematic verification prior to the experiments presented in Figure 2E to ensure that sterol esterification has been effectively prevented by avasimibe. In more detail, Nile Red dyes LD, Filipin dyes free cholesterol. **D.** Cell viability in response to oleate (60 μM), zeocin (10 μg/mL) or combination of both treatments was assessed by crystal violet assay. 30 µM BSA (mock) was used as control for oleate incubation. Briefly, RPE-1 and Huh-7 cells were seeded in 96-well plates, one plate per time point (see Materials and Methods for more details). The indicated treatments were added at day 0 and renewed at day 3. Every 24h, one plate was processed to read Crystal violet absorbance at 570 nm. The absorbance values are proportional to the number of cells and were normalized to day 0 to calculate the proliferation rate. Graphs show the mean and the standard deviation out of three independent experiments. **E.** RPE-1 and Huh-7 cells were incubated with oleate (60 μM) or 30 µM BSA (mock) for 2h, followed by addition of zeocin (10 μg/mL) where indicated for 2 extra hours. BrdU (10 μg/mL) was added to all samples during the last 2 hours of treatment. Cells were fixed and processed for BrdU detection using the BD Pharmingen™ BrdU Flow Kit (Cat. No. 552598) according to the manufacturer’s instructions. DNA was stained with DAPI (20 μg/mL). Cells were analyzed by cytometry and assigned to each phase of the cell cycle (subG1: less than 1C DNA content, G1: 1C DNA content and BrdU-, S: BrdU+, G2: 2C DNA content and BrdU-). Graphs show the mean and the variability of 2 independent experiments.

**Figure S3.**
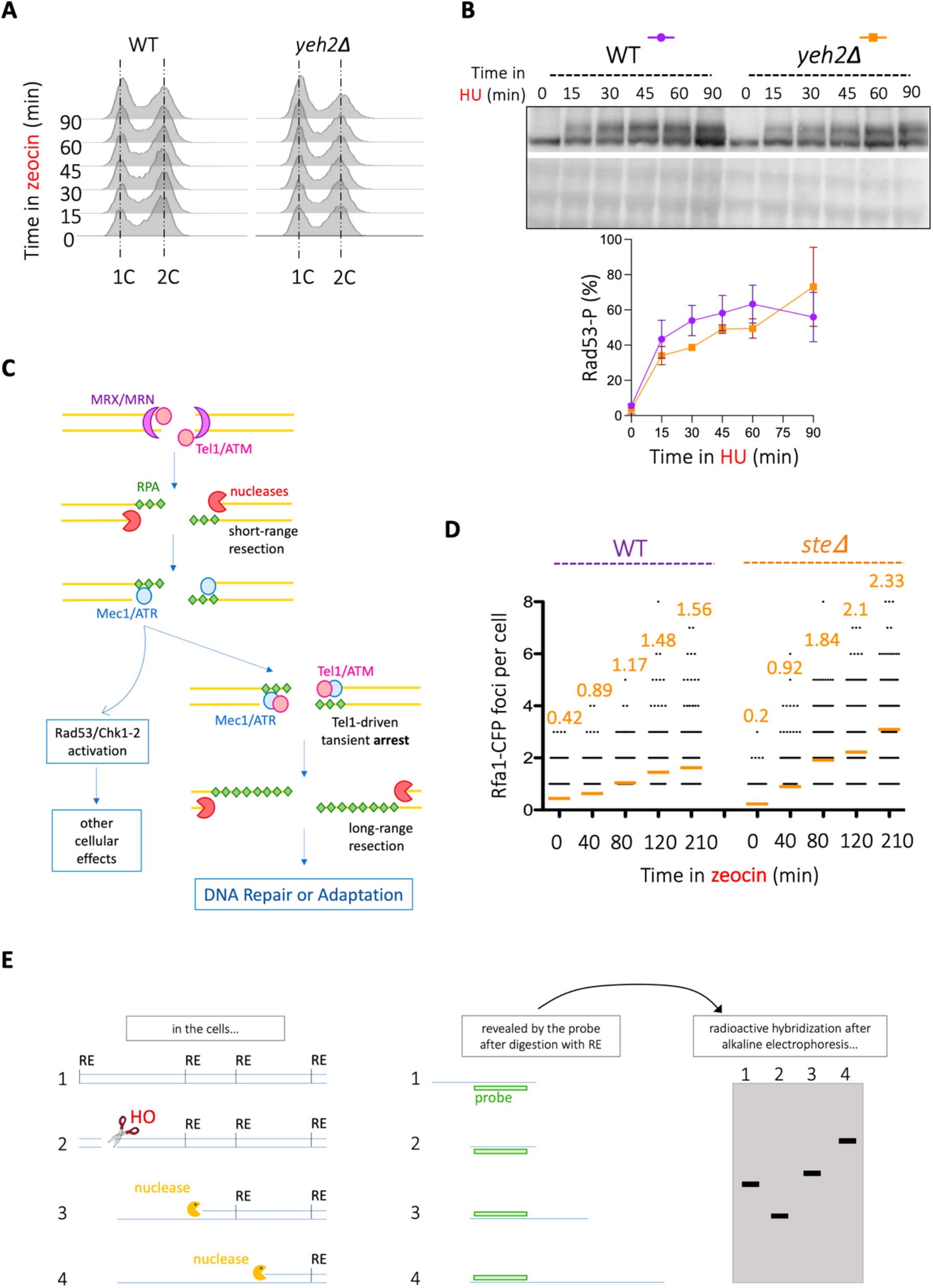
Additional information related to Figure 3. **A.** Cytometry profiles concerning the experiment presented in Figure 3A. **B. Top:** identical experiment as that reported in Figure 3A but in response to 20 mM hydroxyurea (HU). **Bottom:** quantification of the percentage of Rad53 molecules being phosphorylated at each time-point. Details of quantification and plotting as in Figure 3A. **C.** Simplified scheme of the detection, signalling and repair of a DNA DSB: In response to a DSB, the MRX/MRN complex and Tel1/ATM bind the broken tips and allow the subsequent resection of 5’ ends by nucleases. This leads to the formation of ssDNA that becomes coated by RPA, thus priming the recruitment of Mec1/ATR. This kinase phosphorylates on-chromatin targets, which serve to transiently limit the extent of resection, and also effector proteins, as Rad53 and CHK1-CHK2, which coordinate cell status with the need of repair. It is at this stage that Tel1/ATM acts again by triggering a transient arrest before the onset of long-range resection ^41^. Eventually, Tel1/ATM disengages, which permits long-range resection. If a homologous sequence is found, productive repair by homologous recombination (HR) will follow. In the absence of a repair template, the cell can resume cycling even in the absence of repair, a phenomenon known as adaptation. **D.** Exponentially growing *S. cerevisiae* WT and *steΔ* cells were treated with 100 µg/mL zeocin and samples retrieved at the indicated timepoints for visualization by fluorescence microscopy. The establishment of DNA resection factories was assessed by counting the number of Rfa1-CFP foci per cell, which is plotted in this graph. The mean value of each timepoint is indicated by an orange bar and number. This is one illustrative experiment of the 3 that were carried out to build the graph presented in Figure 3B. **E. Left** panel: (1) the double stranded genomic DNA is represented by straight blue lines, where the restriction sites for a given restriction enzyme (RE) are indicated. (2) the addition of galactose to the cultures induces the expression of the HO nuclease, whose target site is cut. (3, 4) resection by nucleases progressively destroys RE sites in the genome. **Middle** panel: the use of a radioactive probe will reveal fragments of different sizes depending on the HO / RE / nuclease profile. **Right** panel: migration of these fragments in an alkaline electrophoresis followed by Southern blot with the indicated probe detects each resection intermediate.

**Figure S4.**
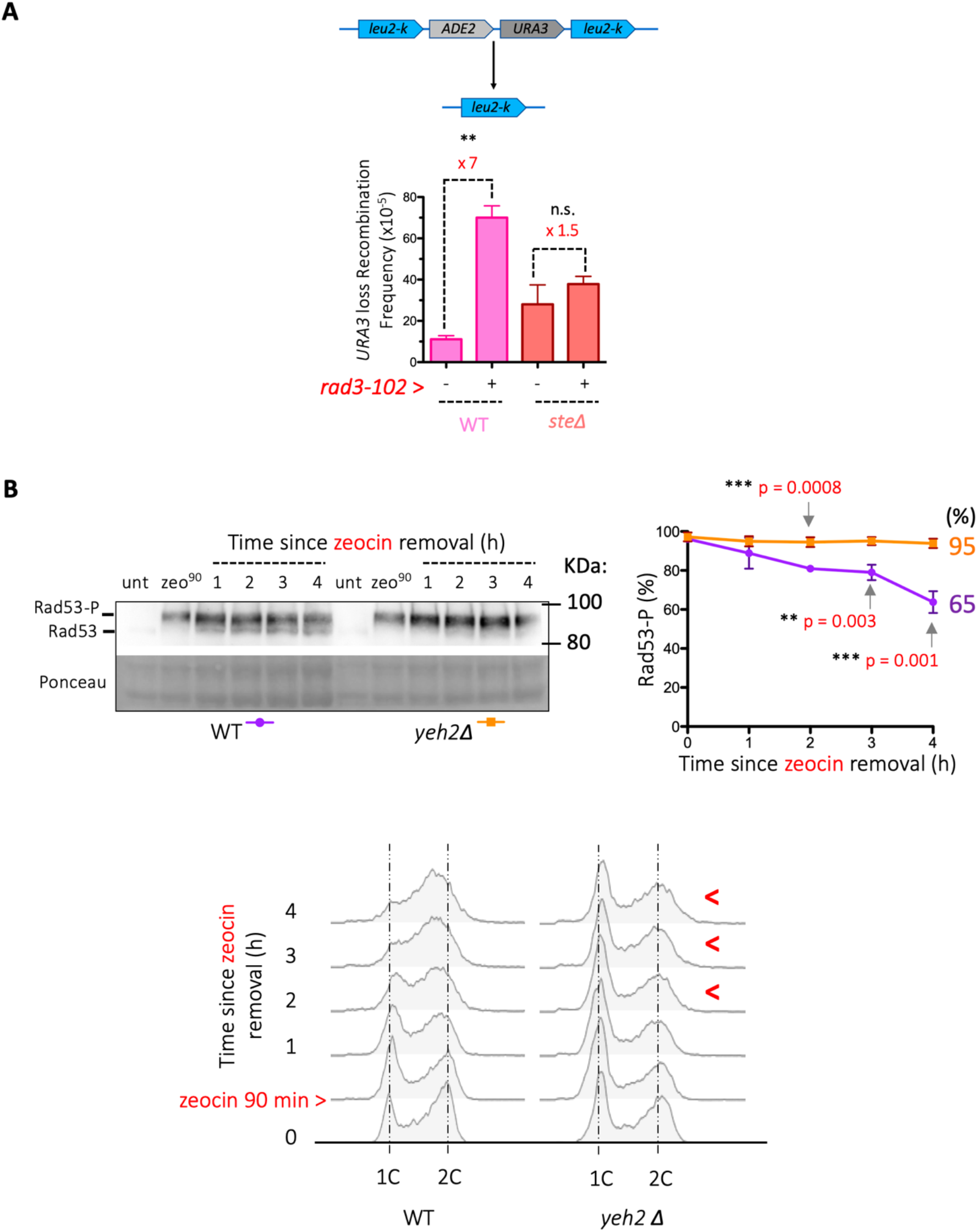
Additional information relating to Figure 4. **A. Top:** scheme of the genome-integrated system allowing the analysis of HR rates, mostly through single-strand annealing. Details as in Figure 4A. **Bottom:** for each recombination test, the 4 strains of interest (WT, *rad3-102*, *steΔ* and *rad3-102 steΔ* cells) were streaked onto YPD plates, and 6 isolated colonies out of each plate used to measure the number of recombinant and total cells, thus yielding the shown recombination frequencies. For every 6 frequencies derived from 6 colonies, one median frequency is calculated (= 1 experiment). Each bar represents the mean frequency of the median of at least 3 independent experiments. *rad3-102*-induced recombination is indicated as a fold-change. Statistical evaluation of the differences between the recombination frequencies was assessed using a *t*-test: n.s., non significant; **, *p*-value < 0.001. **B.** Identical to Figure 4D but to compare the DDR de-activation in WT *versus yeh2Δ* cells.

**Figure S5.**
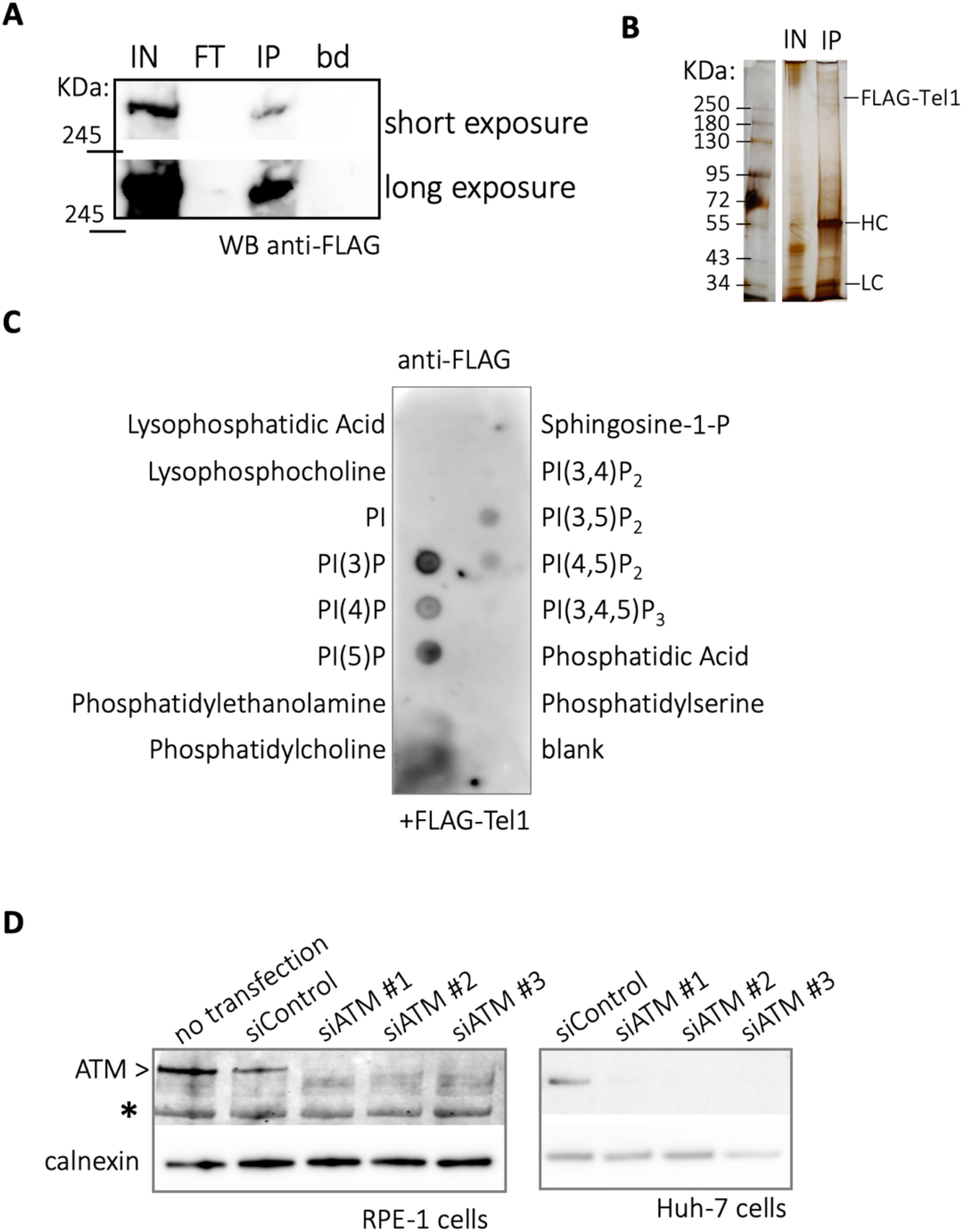
Additional information related to Figure 6. **A.** *S. cerevisiae* cells transformed with a galactose-inducible FLAG-Tel1 construct were grown overnight in YPGal medium, treated for 2 hours with 100 µg/mL zeocin under the assumption that this will promote the release of Tel1 from PI(4)P. FLAG-Tel1 was immunoprecipitated using M2-agarose anti-FLAG beads and retrieved by competitive elution using an excess of 3xFLAG peptide. The displayed Western blot illustrates the extract preparation process. IN: input; FT: flow-through; IP: competitively eluted FLAG-Tel1; bd: Laemmli buffer + boiling-eluted remnants on the beads. One sixth of the eluted Flag-Tel1 was loaded as “IP”. All the Laemmli buffer volume used to boil the beads was loaded. **B.** Cells treated as in (A) were used to prepare GFP-Tel1. A sample of the total extract is loaded as the input (IN). FLAG-Tel1 was immunoprecipitated using M2-agarose anti-FLAG beads and released by boiling the beads in Laemmli buffer, the goal being to check as stringently as possible the purity of the preparation (IP). HC, heavy chain of IgG1, expected at 56 kDa; LC, light chain of IgG1, expected at 25 kDa. **C.** A membrane on which 100 pmol of the indicated lipid species were spotted was incubated with immunopurified FLAG-Tel1 and further developed using an anti-FLAG antibody. The membrane shown is illustrative of one out of four independent experiments. * indicates one unspecific band. **D.** Western blot to detect total ATM levels (calnexin used as a loading control) for the experiments shown in Figure 6C bottom right, in which cells were transfected with one siRNA control or three independent siRNAs against ATM, one at a time, prior to performing PLA.

**Figure S6.**
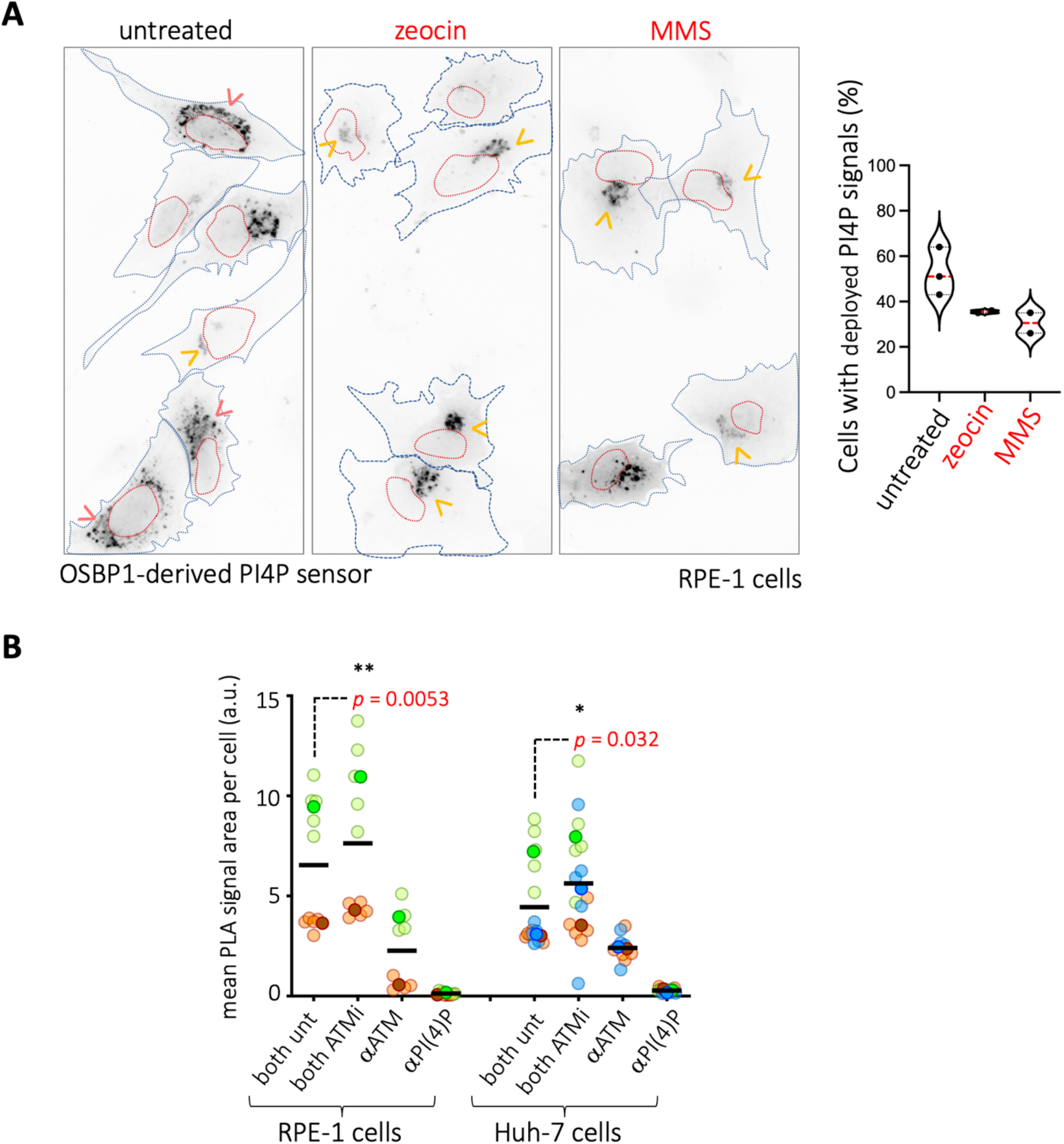
Additional information for Figure 7. **A.** RPE-1 cells were transfected with a vector expressing a sensor capable of revealing PI(4)P. The sensor is composed by the domain from OSBP1 that binds PI(4)P followed by a GFP moiety^79^. The cells were either left untreated or exposed to 0.005% MMS or to 10 µg/mL zeocin for 2 h. Upon fixation, DNA was stained using DAPI. The images shown were acquired on the GFP channel to observe PI(4)P signals. DAPI signals were used to draw the nuclei contour and impose it on the GFP ones (red dashed lines). The cell boundaries were revealed by forcing the contrast and underlined using blue dashed lines. Extended PI(4)P signals are indicated by pink arrowheads, while compact PI(4)P signals are marked by yellow ones. The graph on the right in which the percentage of cells displaying PI(4)P signals is quantified for three independent experiments. The median value of each violin plot is shown by a dashed red line. **B.** Proximity ligation assay (PLA) to assess whether ATM and PI(4)P proximity is altered when cells are pre-treated for 1 h with 500 nM of the ATM inhibitor (ATMi) AZD0156. Each point is the value of having measured all the PLA signals present in one photo (approximately 20 to 40 cells) and divided this value by the number of nuclei. Each independent experiment is plotted in a different colour. Further, the mean value of each independent experiment is highlighted by a more intense colour than the individual values of that experiment, for which the colour is more translucid. Last, the solid horizontal line marks the mean of the means. A one-way ANOVA for multiple comparisons was applied to evaluate whether there were significant differences between the means of the indicated conditions.

**Figure S7.**
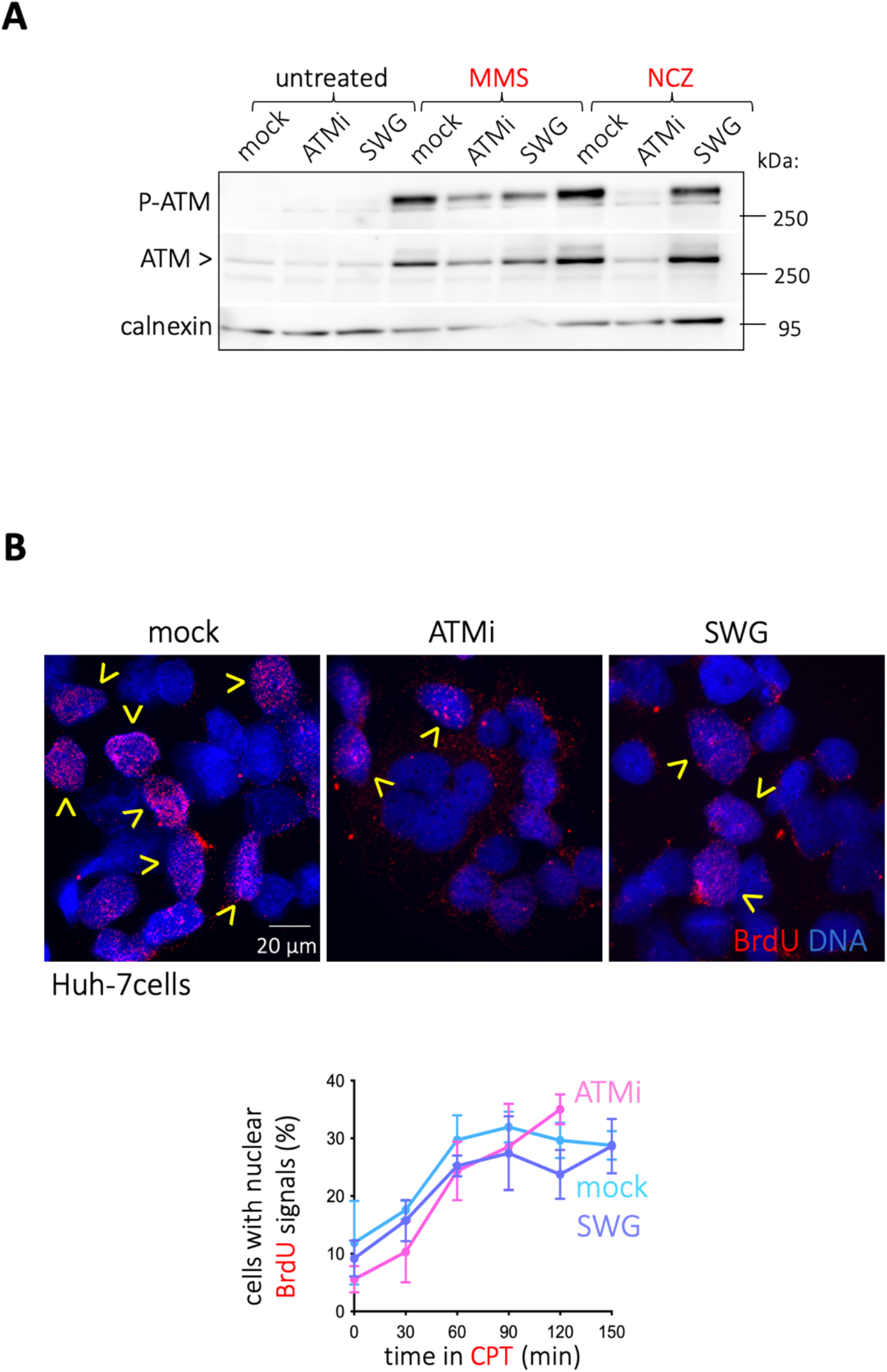
Additional information for Figure 8. **A.** Example of an experiment as the one shown in Figure 8A that also includes the detection of total ATM. As observed, the overall stability of the protein also seems to be impacted by the treatments. This is why calnexin is used as a referent for the load. **B.** Huh-7 cells were grown with 10 µg/mL BrdU present in the culture medium for the last 16 h prior to further processing. Cells were then either mock-treated, or pre-treated for 1 h with 500 nM of the ATM inhibitor (ATMi) AZD0156 or 10 nM of the specific OSBP1 inhibitor schweinfurthin G (SWG). Then, 200 µM CPT (see Materials & Methods) was added and cells were processed for BrdU detection (see M&M) at the indicated times. **Top:** illustrative images of BrdU-positive nuclei, indicated by yellow arrowheads. **Bottom:** the percentage of cells in the population whose nuclei were positive for BrdU was established by visual inspection of the acquired images. The graph shows the mean percentage out of three independent experiments and the error bars represent their associated SEMs.

## Materials and Methods

### Reagents

filipin (F4767, Sigma-Aldrich), avasimibe (PZ0190, Sigma-Aldrich), Nile Red (HY-D0718, CliniSciences), 5-Fluoroorotic acid monohydrate (5-FOA, F595000, Toronto Research), oleic acid for *S. cerevisiae* cells (O1008, Sigma-Aldrich); oleic acid:BSA conjugate for human cells (O3008, Sigma-Aldrich), Hoechst (B2261, Sigma-Aldrich), methyl metanosulfonate (129925, Sigma-Aldrich), zeocin (R25001, ThermoFisher), nocodazole (M1404, Sigma-Aldrich), hydroxyurea (H8627, Sigma-Aldrich), camptothecin (C9911, Sigma-Aldrich), 3xFLAG peptide (F4799, Sigma-Aldrich), Fatty Acid-free BSA (A6003, Sigma-Aldrich), DAPI (D9542, Sigma-Aldrich), cOmplete protease inhibitor cocktail (11836170001, Roche), Halt Phosphatase inhibitor cocktail (78420 ThermoFisher) and ProLong (P36930, ThermoFisher), BrdU (5-Bromo-2ʹ-deoxyuridine, B5002-1G, Sigma-Aldrich), AZD0156 (HY-100016-5mg, Clinisciences), neocarzinostatin (NCZ, N9162-100UG, Sigma-Aldrich), schweinfurthin G (SWG) was a kind gift from Bruno Mesmin and Bruno Antonny, Crystal violet (V5265-250ML, Sigma-Aldrich). siControl was the universal negative control siRNA #1 (reference SIC001, Sigma-Aldrich), and the siRNAs against ATM were also from Sigma-Aldrich: siATM#1(SASI_Hs01_00093615, 5’-CUUAGCAGGAGGUGUAAAU[dT][dT]-3’); siATM#2 (SASI_Hs01_00093616, 5’-CCCAUUACUAGACUACGAA[dT][dT]-3’); siATM#3 (SASI_Hs01_00093617, 5’-GUUACAACCCAUUACUAGA[dT][dT]-3’).

### Antibodies

anti-calnexin (610523, BD transduction Laboratories, WB at 1/1000), anti-Rad53 (gift from C. Santocanale, WB at 1/3000), anti-P-Thr68-CHK2 (2661S, Cell Signaling, WB at 1/1000), anti-total CHK2 (05-649, Millipore, WB at 1/1000), anti-total ATM (GTX132147, GenTex, IF at 1/200, PLA at 1/2000), anti-P-Ser1981-ATM (4526, Cell Signaling, IF at 1/200, WB at 1/1000), anti-FLAG (F3165, Sigma-Aldrich, WB at 1/3000), anti-FLAG^R^ M2 affinity gel (A2220, Sigma-Aldrich), anti-ssDNA (Enzo; ALX-804-192-R200, used at 1/500), anti-PI4P (Z-P004; Echelon Biosciences, PLA at 1/2000), anti-BrdU (Purified Mouse Anti-BrdU Clone 3D4, 15838318, FisherScientific, IF at 1/100) and anti-tubulin (T5168, Sigmal-Aldrich, WB at 1/3000). Secondary antibodies for WB were goat anti-rabbit (A0545, Sigma-Aldrich) at 1/5000 and rabbit anti-mouse (A9044, Sigma-Aldrich) at 1/5000; for IF DyLight 488 donkey anti-rabbit (406404, Biolegend) at 1/500 and DyLight 649 goat anti-mouse (405312, Biolegend) at 1/500.

### *S. cerevisiae* cells culture and treatments

Cells were grown at 25°C in YEP medium supplemented with 2% glucose unless otherwise indicated. In experiments where a plasmid needed to be maintained, the cells were grown overnight in YNB lacking the relevant aminoacid for selection, then diluted in the morning and left to reach exponential phase in YEP medium supplemented with 2% glucose for approximately 4 hours. For expression of constructs under the control of the *GAL1* promoter, cells were grown overnight in the appropriate medium + 2% Raffinose, then induction was started by adding 2% Galactose. All experiments were performed with asynchronous cultures of cells growing exponentially. For liquid culture time-course experiments, final concentrations of the drugs were: 100 µg/mL zeocin, 0.1 M HU, 100 µM CPT, 15 µg/mL nocodazole. The pre-treatment with oleate in liquid cultures was done using a 0.05% concentration for 2 hours. Strains used in this study are either shown in Table S1, or derived from crosses. Additionally, endogenous *PUS1* was N-terminally tagged with mCherry by *Bgl*II-linearising then transforming YIplac211-mCherry-*PUS1* ^80^. Plasmids used in this study are shown in Table S2.

### Human cell culture and treatments

RPE-1 or Huh-7 were seeded on p100 plates (dilution 1:7 from one confluent p100). After two days, medium was changed (at 37 °C) and treatments were applied. RPE-1 were grown in DMEM (D5796-500ML, Sigma-Aldrich) and Huh-7 in DMEM+GlutaMAX (Gibco, 31966-021) supplemented with 10% FBS (S1810-500, Biowest) and 1% Pen/Strep (P0781, Sigma-Aldrich). At the end of the corresponding treatments, cells were collected for Western blot or fixed for immunofluorescence with 4% PFA/PBS (20 min at room temperature) or cold methanol in the case of P-ATM experiments (chilled at –20°C; fixation for 10 min), and then washed once with PBS. **Transfections:** Cells were seeded 24 hours prior to transfection with XtremeGENE 360 (08724121001, Merck). Following the supplier’s protocol, for plasmid transfection, 2µL of the reagent were added per for 1µg of DNA in a 100 µL solution. For siRNA transfection, 7.5 µL of the reagent were combined with 40 nM siRNA in 6-wells-plate wells and left 28 hours or 72 hours, respectively, prior to the intended experiment. **Treatments:** DNA-damaging treatments were always applied for 2 hours at a final concentration of 0.005% MMS, 10 µg/mL zeocin, 120 ng/mL neocarzinostatin (NCZ). Also, due to a calculation mistake, we did our experiments with 200 µM camptothecin (CPT), which is a 100-fold higher dose than usually seen in the literature. Yet, at this high dose, cells were fit in the time frame of our experiments, probably because once CPT has trapped all topoisomerase-I molecules in the cell, further addition of the drug does not have any additional impact. We have repeated once more the experiments presented in Figures 8A, 8B and S7B with 2 µM CPT and we have confirmed that the conclusions do not change. Pre-treatments were done for 4 hours with 60 µM oleate, or 2 hours with 5 µM avasimibe.

### BrdU immunofluorescence detection

Cells were seeded on coverslips at a 50-60% confluency and the next day BrdU was directly added at a final concentration of 10 µg/mL and incubated for 16h. Cells were then pre-treated with 500 nM AZD0156 for 1h or 10 nM SWG for 2h, or mock-treated, then exposed to 200 µM CPT and coverslips retrieved at 0, 30, 60, 90, 120 or 150 minutes. Cells were then pre-extracted, fixed and immunostained exactly as previously described ^56^.

### Dot Blot to detect ssDNA

RPE-1 cells were seeded in p100 plates to reach 90% confluency the next day, then were incubated with 60 µM oleate-(BSA) or 30 µM BSA (mock) for 2 hours. 0.005% MMS was added, and cells were trypsinized and collected at the indicated times. Cell pellets were stored at −20°C. To extract genomic DNA (gDNA), cell pellets were incubated with lysis buffer (10 mM Tris pH 8; 1mM EDTA; 0.5% SDS; 60 µg/mL Proteinase K (EU0090-C, Euromedex) overnight at 37°C. 1 volume of Phenol:Chloroform:Isoamyl Alcohol (25:24:1) (P2069-400ML, Sigma-Aldrich) was added to the lysates, then transferred to Phase Lock Gel Tubes (733-2477, VWR) and centrifuged at 12000 g for 5 min. The top layer was retrieved and incubated with 1 mg/mL (11508686, FisherScientific) 1 h at 37°C. gDNA phase was isolated using again 1 volume of Phenol:Chloroform:Isoamyl Alcohol (25:24:1) in Phase Lock Gel Tubes and centrifuged at 12000 g for 5 min. gDNA was precipitated using 0.3M NaOAc pH 5 and 2.4 volumes of ethanol 100%, and centrifugation at 17000 g for 15 min. gDNA was washed with 70% ethanol, and resuspended in TE buffer (10 mM Tris pH 8; 1mM EDTA). A Hybond-N+ membrane (GERPN203B, Sigma-Aldrich) was pre-hydrated with 6x SSC (0.9 M NaCl; 90 mM Trisodium citrate), and wells were washed with TE buffer. 1.5 µg of gDNA were dissolved in 250 µL TE buffer and loaded onto the membrane with the help of a Bio-Dot apparatus (Bio-Rad, 1706545) and a vacuum-pump, then washed with 2X SSC. DNA was crosslinked to the membrane using 12 µJoules/cm^2^ UV-C. The membrane was next blocked with 3%BSA/TBS-T for 1 h, incubated with anti-ssDNA at 1:500 in TBS-T/1% BSA/0.01% sodium azide overnight at 4°C, washed and further incubated with anti-mouse antibody at 1:5000 in 5% milk/TBS-T for 45 min, then developed using SuperSignal^TM^ West Pico PLUS (ThermoScientific).

### Proximity Ligation Assay

Cells were seeded on coverslips and, after the pertinent treatment, fixed with 4% PFA/PBS during 20 min at room temperature. Subsequently, every step took place in a humid chamber using the PLA reagents from Sigma-Aldrich. Cells were permeabilized with 0.5% Triton/PBS for 10 min and incubated with Blocking Solution from the Duolink *in situ* PLA kit (DUO92013) for 1 h. Primary antibodies (anti-PI(4)P and anti-ATM) were diluted at 1:2000 in the Antibodies Diluent from the kit and incubated overnight at 4°C onto the coverslips. For technical controls, one of these primary antibodies was omitted at a time. PLA minus and plus probes (DUO92004 and DUO92002, Sigma-Aldrich) were next mixed and incubated in the Blocking Solution for 20 min as specified by the supplier, then added onto the coverslips and incubated 1h at 37°C. Coverslips were washed twice with Buffer A (150 mM NaCl; 10 mM Tris; 0.05% Tween-20; pH 7.4), then incubated with Ligation Mix (1:40 ligase; 1:5 ligase buffer) 30 min at 37°C, washed again twice with Buffer A, then incubated with Polymerization Mix (1:80 polymerase; 1:5 polymerase buffer) 1h at 37°C. Last, coverslips were washed with Buffer B (100 mM NaCl; 200 mM Tris; pH 7.5), then with 0.01X Buffer B. The coverslips were finally mounted with Duolink *in situ* Mounting Medium with DAPI (DUO82040), then sealed with nail polish.

### Crystal violet proliferation assay

RPE1 and Huh-7 cells were seeded in 96-well plates at an initial number of 1000, 2000 or 4000 cells per well (4 wells per initial seeding number). One independent plate was seeded per time point. Once cells were adhered, the specified treatments were applied and incubated for 1 to 6 days. Medium was changed and all treatments were renewed at day 3. Every 24h, one plate was washed twice with PBS, fixed with 50 µL of 4% PFA-PBS for 10 min at room temperature, stained with 80 µL of 0.1% crystal violet in water for 30 min at room temperature, and extensively washed with distilled water to eliminate the excess of dye. The water was removed and cells were allowed to dry before solubilization with 200 µL of 10% acetic acid. Absorbance was read at 570 nm with a Multiskan microplate reader (ThermoFisher). Crystal violet absorbance directly correlates with the number of living cells that remain attached to the plate. For Day 0, one extra plate was seeded at the same time than the others but no treatments were added. This plate was fixed after 7 hours following the same protocol to assess the initial number of cells per condition. Absorbance at 570 nm at each timepoint was normalized to the values from day 0 in order to calculate the proliferation rate.

### Telomere length

was measured by PCR after end labeling with terminal transferase ^53, 54^. End-labeling reactions (40 μL) contained 120 ng genomic DNA, x1 New England Biolabs™ Terminal Transferase Buffer, 1 mM dCTP, 4 units Terminal Transferase (New England Biolabs™) and were carried out at 37°C for 30 minutes followed by heat inactivation at 75°C for 10 minutes. 1/5^th^ volume of 5 M NaCl, 1/80^th^ volume of 1 M MgCl₂ and 1 volume of isopropanol were added to the reaction and DNA was precipitated by centrifugation at 17000g during 15 min. Precipitated DNA was resuspended in 40 μL of ddH₂O. The end-labeled molecules were amplified by PCR using the primer 5’-GCGGATCCGGGGGGGGGGGGGGGGGG-3’ and 5’-TGTGGTGGTGGGATTAGAGTGGTAG-3’ (X) and 5’-TTAGGGCTATGTAGAAGTGCTG-3’ (Y’), respectively. PCR reactions (50 μL) contained between 40 ng and 80 ng of DNA, 1x myTaq buffer, and primers 0.4 μM each. Amplification was carried out with 5 U of MyTaq polymerase (BIO-21105, Meridian Biosciences). The conditions were 95°C, 5 minutes; followed by 35 cycles of 95°C, 1 minute; 56°C (Y reaction) / 60°C (X reaction), 20 seconds; 72°C, 5 minutes. Reaction was ended with 5 minutes at 72°C. Samples were visualized in a 2 % agarose gel containing 1× GelRed (BTM41003, Ozyme).

### Immunofluorescence of human cells

cells fixed on coverslips were washed once with PBS and permeabilized with 0.2% Triton/1x PBS for 10 min at room temperature, then saturated with 3% BSA/1x PBS for 30 min. Coverslips were incubated with primary antibodies diluted in 3% BSA/1x PBS for 90 min then washed 3 times x 10 min with 1x PBS under gentle shaking. Coverslips were further incubated with secondary antibodies diluted (1:500) in 3% BSA/1x PBS for 45 min while protecting them from light from this point on, then washed 3 x 10 min with 1x PBS under gentle shaking, then incubated for 5 min with Hoechst (20 µg/mL) diluted in H_2_O, and washed 3 times with H_2_O. Finally, coverslips were allowed to dry and mounted using ProLong, then left to dry overnight at room temperature in the darkness.

### Western blot from human cells

all the samples were lysed with High Salt Buffer (50 mM Tris pH 7.5, 300 mM NaCl, 1% Triton X-100, Protease inhibitors (Roche), Phosphatase inhibitors Halt cocktail) (400 µL per p100) for 10 min on ice with frequent vortexing. Samples were centrifuged 10 min at 14000 rpm at 4°C, and supernatants quantified using the Pierce^TM^ BCA kit (10741395, ThermoFisher). For Western Blot, 20-30 µg of whole cell extracts were loaded onto home-made 10% acrylamide gels (1.5 mm thick) and migrated 70 minutes at 40 mA per gel (migration buffer: 25 mM Tris, 200 mM Glycine, 0.1% SDS). The proteins were transferred to a nitrocellulose membrane for 2 hours at 100 V in transfer buffer (25 mM Tris, 200 mM Glycine, 20% ethanol). For detection of P-ATM, whole cell extracts were loaded on 3-8% acrylamide gradient gels (1.5 mm-thick) and migrated 90 min at 40 mA (migration buffer: 50 mM Tris, 50 mM Tricine, 0.1% SDS). The proteins were transferred to a nitrocellulose membrane overnight at 30 V in transfer buffer (25 mM Tris, 200 mM Glycine, 20% methanol).

### FLAG-Tel1 immunoprecipitation

A 250 mL-culture of cells bearing the p*GAL1p*-FLAG-*TEL1* vector reaching a density of 7×10^6^ cells/mL in YEP + 2% Galactose was treated for 2h with 100 µg/mL zeocin at 25°C. The culture was centrifuged at 4000 g for 3 minutes and the supernatant was discarded. The cell pellet was washed once with Disruption Buffer (20mM Tris-HCl pH7.9, 10mM MgCl_2_, 1mM EDTA, 10% glycerol, 0.3M ammonium sulfate and protease inhibitor). The pellet was subsequently resuspended in 1 volume of Disruption Buffer and 2 volumes of pre-cooled Glass Beads (0.4 mm diameter) then subjected to 10 cycles of 30 s Cellbreaker/30 s on ice. Each sample was centrifuged briefly, supernatants were retrieved and further centrifuged for 1h at 12000 g at 4°C. Supernatants were collected again and protein concentration estimated by the Bradford method. Next, 40 µL of M2-agarose beads were centrifuged at 5000 g during 30 s, supernatant was discarded then M2-beads were washed two times with 500 µL of 1x TBS. 300 µg of protein extract were added to the beads and diluted up to 1 mL using 1x TBS + protease inhibitor cocktail. The mixture was incubated at 4°C overnight on a rotating wheel. Next morning, the mixture was centrifuged at 5000 g during 30 s, supernatant was removed (and saved as the “flowthrough fraction”) and the beads were washed three times with 500 µL 1x TBS. 250 µL of 3XFLAG peptide (stock at 1 mg/mL) were added and incubated 30 minutes at 4°C on a rotating wheel. The mixture was centrifuged 30 s at 5000 g and the supernatant recovered and used for lipid strip incubation and for Western Blot verification. The beads were boiled 5 min using 30 µL of 1X Laemmli Buffer. These 30 µL were used to evaluate by Western Blot the eventual, residual presence of FLAG-Tel1 onto the beads. For silver staining of proteins, samples were migrated in a 7.5% acrylamide gel (Bio-RAD, 4568023, USA) at 150V for 45 min. Then, the gel was treated with and according to SERVA Silver Staining kit for SDS PAGE (SERVA Electrophoresis GmbH, 35077, Germany), performing the fixation step overnight and the developing one for 5 min.

### Western blot from *S. cerevisiae* cells

For the regular detection of Rad53, approximately 5 × 10^8^ cells were collected at each relevant time point and washed with 20% trichloroacetic acid to prevent proteolysis, then resuspended in 200 µL of 20% trichloroacetic acid at 4°C. The same volume of glass beads was added, and cells were disrupted by vortexing for 10 min. The resulting extract was spun for 10 min at 1000 g also at room temperature and the resulting pellet resuspended in 200 µL of Laemmli buffer. Whenever the resulting extract was yellow-colored, the minimum necessary volume of 1 M Tris base (non-corrected pH) was added till blue color was restored. Then, water was added till a final volume of 300 µL was reached. These extracts were boiled for 10 min and clarified by centrifugation as before; 10–15 µL of this supernatant was loaded onto a commercial 3–8% acrylamide gradient gel (BioRad) and migrated 70 min at 150 V to separate Rad53 isoforms, then proteins transferred to a nitrocellulose membrane. Detection by immunoblotting was performed with anti-Rad53 antibody, a kind gift from Dr. C. Santocanale, Galway, Ireland. To detect FLAG-Tel1, samples were loaded onto a commercial 7.5 % acrylamide gradient gel (BioRad) and migrated 60 min at 60 mA, then proteins transferred to a nitrocellulose membrane over night at 4 °C.

### Lipids membrane hybridization

A commercially available membrane on which 100 pmol of each of the lipids of interest had been spotted (P-6002 or P-6001, Echelon Biosciences) was blocked using TBS-Tween_0.1%_ supplemented with 3% fatty acids-free BSA during 1h in the dark at room temperature. 5/6^th^ of the M2-immunoprecipitated FLAG-Tel1 preparation were incubated on the lipid membrane in a final volume of up to 1 mL (TBS-Tween_0.1%_ supplemented with proteases inhibitor) sealed in a plastic bag, then incubated 4°C in the dark under soft shaking. The following day, the lipid membrane was washed three times in TBS-Tween_0.1%_ (10 minutes each wash) and the anti-FLAG antibody incubated during 3h at 4°C in the dark. All subsequent steps were performed as for regular Western Blot excepted that all incubations were done in the dark.

### Recombination analyses

The genome-integrated system *leu2-k::ADE2::URA3::leu2-k* ^39^ allows the analysis of the frequency of homologous recombination, mostly that occurring through single-strand annealing. The loss of the *URA3* marker, which can be selected for in plates containing 5-FOA (0.5 g/L) permits the calculation of the recombination frequency. For each recombination test, cells were streaked onto plates containing either 7.5 µg/mL zeocin or not, and either 0.05% oleate or not, as indicated, and incubated for 3 days at 26°C. From each plate, 6 isolated colonies were resuspended in 1 mL sterile water each, dilutions of each tube done and dilutions from 1/4^th^ to 1/400^th^ seeded on 5-FOA-containing plates in order to select for recombinants, and one dilution 1/40000^th^ seeded on YPD plates to count the total number of cells, finally allowing to calculate the recombination frequencies. For every 6 frequencies derived from 6 colonies, one median frequency was calculated (= 1 experiment). The displayed results for each condition represent the mean frequency out of the median of at least 3 independent experiments.

### Pulsed Field Gel Electrophoresis

Human cells in culture treated as indicated were retrieved in 1x PBS and counted. Approximately 6×10^5^ cells were used per agarose plug. Typically, each plug was created by mixing very smoothly 50 µL of this cell suspension and 50 µL of low melting point agarose pre-prepared at 1 % in 1x PBS, and pouring the mix into a disposable (yet re-used) plug mold (Bio-Rad Laboratories). Plugs were allowed to solidify for 30 min at room temperature and 30 min at 4°C. They were then incubated (typically 5 plugs of each condition in 2 mL of buffer) in lysis buffer (100 mM EDTA, 1% [wt/vol] sodium lauryl sarcosine, 0.2% [wt/vol] sodium deoxycholate, and 1 mg/mL proteinase K) at 37°C for 24 h. Plugs were then washed five times in 5 mL of 20 mM Tris-HCl, pH 8.0, 50 mM EDTA, and were then ready to be loaded onto an agarose gel. Electrophoresis was performed for 21 h at 14°C in 0.9% (wt/vol) Pulse Field Certified Agarose (Bio-Rad Laboratories) in exactly 2.4L of 0.5x Tris-borate/EDTA buffer in a Rotaphor apparatus (Biometra, Analitik Jena) with the following protocol: block I: 9 h, 120° included angle, 5.5 V/cm, 30 to 18-s switch; block II: 6 h, 117° included angle, 4.5 V/cm, 18 to 9-s switch; block III: 6 h, 112° included angle, 4.0 V/cm, 9 to 5-s switch. The gel was then stained with ethidium bromide, photographed and well and in-gel signals analyzed using Image J. Relative DSB levels were calculated by dividing the in-gel signal between the total lane signal and expressed as a percentage.

### Analysis of DNA content by flow cytometry in *S. cerevisiae*

430 µL of culture samples at 10^7^ cells/mL were diluted in 1 mL of 100% ethanol. Cells were centrifuged for 1 minute at 16000 g and resuspended in 50 mM Na-Citrate buffer containing 5 µL of RNase A (10 mg/mL, Euromedex, RB0474) for 2 hours at 50°C. 6 µL of Proteinase K (Euromedex, EU0090-C) were added for 1 hour at 50°C. Cell aggregates of cells were dissociated by sonication (one 3s-pulse at 50% potency in a Vibracell 72405 Sonicator). 20 µL of this cell suspension were incubated with 200 µL of 50 mM Na-Citrate buffer containing 4 µg/mL Propidium Iodide (Fisher scientific). Data were acquired and analyzed using a Novocyte Express (Novocyte). At least three independent biological replicates, each counting at least 10000 cells per sample and condition, were analyzed.

### Cytometry for human cells

RPE-1 and Huh-7 cells were mock-treated (30 µM BSA) or treated with oleate (60 μM) for 2h, followed by addition or not of zeocin (10 μg/mL) for 2 extra hours. BrdU (10 μg/mL) was added to all samples during the last 2 hours of treatment. One extra well of mock-treated cells without BrdU was added and processed as the rest of the samples to be used as negative control for FACS analysis. BrdU detection was performed using the BD Pharmingen™ BrdU Flow Kit (Cat. No. 552598) according to the manufacturer’s instructions. More in detail, cells were trypsinized, counted and 0.5 million cells per condition were washed once with 1 mL PBD, subsequently fixed and permeabilized with 100 μL BD Cytofix/Cytoperm Buffer for 15 min at 4°C, then washed with 1 ml of 1X BD Perm/Wash Buffer, further permeabilized with 100 µL of BD Cytoperm Permeabilization Buffer Plus for 10 min at 4°C, washed with 1 mL of 1X BD Perm/Wash Buffer, re-fixed again with 100 μL BD Cytofix/Cytoperm Buffer for 10 min at 4°C, washed with 1 mL of 1X BD Perm/Wash Buffer, resuspended in 100 µL of DNase (diluted to 300 µg/mL in PBS) for 1 h at 37°C, washed with 1 mL of 1X BD Perm/Wash Buffer, incubated with APC tagged anti-BrdU diluted in BD Perm/Wash buffer (1:50) for 20 min at RT, washed with 1 mL of 1X BD Perm/Wash Buffer, DNA was stained with DAPI (20 μg/mL diluted in BD Perm/Wash Buffer) for 10 min at 4°C, washed with 1 mL of 1X BD Perm/Wash Buffer, and resuspended in 250 μL of PBS. Cells were then analyzed by cytometry and assigned to each phase of the cell cycle (subG1: less than 1C DNA content, G1: 1C DNA content and BrdU-, S: BrdU+, G2: 2C DNA content and BrdU-). Graphs show the mean of 2 independent experiments.

### DSB end resection assay

DSB end resection was analyzed in JKM139 (provided by J. Haber, Brandeis University, Waltham, USA)-derivative strains. Overnight mid-log cultures in YEP medium containing 2% raffinose at 1×10^7^ cells/mL were supplemented with 2% galactose to induce HO-dependent cleavage. 50 mL of cells were recovered at each time point and genomic DNA was extracted from cell pellets by vortexing with acid-washed glass beads in a solution containing 200 µL of phenol:chloroform:isoamyl alcohol (25:24:1) and 200 µL of the extraction buffer containing 2% Triton X-100, 1% SDS, 100 mM NaCl, 10 mM Tris pH 8.0, 1 mM EDTA. The DNA-containing aqueous phase was recovered and precipitated with ethanol. Purified DNA was treated with RNAse A and digested with *Ssp*I restriction enzyme over night at 37°C. DNA fragments were then separated by urea-agarose gel electrophoresis in denaturing conditions for 10 hours at 150V. The gel was subsequently transferred to a GeneScreen Plus membrane (Perkin Elmer), which was hybridized with a radioactive probe corresponding to the distal side of the HO cut. We generated the probe by PCR using the following primers HO_probe_JKM139_F 5’-CCCTGGTTTTGGTTTTGTAGAGTGG-3’ and HO_probe_JKM139_R 5’-GAAACACCAAGGGAGAGAAGAC-3’. Radioactive signals were detected using a PhosphorImager (Typhoon IP, GE) and quantitative analysis of DSB resection was performed with the Image J software by calculating the ratio of band intensities for ssDNA relative to the HO-cut band. Three independent biological replicates were performed.

### Microscopy analyses and image acquisition

Images of fixed human cells were acquired using an ApoTome-equipped microscope (Zeiss). ApoTome technology allows the acquisition of optical sections free of scattered light in order to analyze one focal plane, which allows higher resolution than wide-field microscopy. Filipin staining was achieved by incubating previously PFA-fixed cells in PBS with 25 µg/mL filipin for 2 hours, at room temperature and in the dark, followed by PBS washes. Detection was carried out in the UV range (360/460nm). For *S. cerevisiae* cells, 1 mL of the culture of interest was centrifuged at each relevant time-point of the experiment, the excess supernatant thrown away and the pellet resuspended in the remaining ≈ 50 µL. 3 µL of this cell suspension were directly mounted on a coverslip for immediate imaging of the pertinent fluorescently-tagged protein signals. For visualizing LD in human cells, Nile Red was added to the living cells at a final concentration of 0.5 µg/mL during the last 10 minutes prior to fixation. To dye LD in *S. cerevisiae* cells using Nile Red, 1 µL of a 1 mg/mL stock was added and mixed to the ≈ 50 µL centrifuged pellet (with residual medium) prior to mounting. 3 µL of living *S. cerevisiae* cells were deposited between a coverslip and a slide and directly imaged. The imaging of living *S. cerevisiae* cells was achieved using a Zeiss Axioimager Z2 microscope controlled either by ZEN or by Metamorph softwares.

### Quantifications, plots and statistical analyses

Quantification of the number of LD *per S. cerevisiae* cell, of repair foci per cell, of the percentage of cells in the population displaying foci, BrdU-positive cells, or deployed OSBP1 biosensor signals were achieved by visual inspection and counting by the experimenter. Quantification of the number of P-ATM-positive cells was achieved by a combined Image J macro-assisted detection of positive signals above a given threshold, and further validation and manual counting of positive cells by the experimenter. Proximity Ligation Assay signals were detected using an Image J macro subtracting the background by rolling 30 pixels, setting the threshold level between 700 and 65535, applying “Watershed” to the mask, and then analyzing particles whose size is between 0 and 30000 pixels². The integrated density from those signals per image was retrieved and divided by the number of nuclei in the picture, thus yielding the mean integrated PLA signal per image. Each value obtained this way was used to plot a single dot in the graphs shown in the experiments. Raw signals from membranes (ssDNA, P-ATM, P-CHK2, Rad53, FLAG, PFGE-derived ethidium bromide signals) were quantified for their raw intensity using non-saturated tif images and the “Analyze > Gel” tool from the Image J software and directly plotted without any manipulation. To create the in-gel molecules signal plots presented in Figure 4B,C a line was drawn through the signals in Image J, the pixel intensity along the line drawn with the command “k”, and the associated values exported. GraphPad Prism (version 5 or version 9) was used to plot all the graphs and to statistically analyze the data.

---

**Supplementary Table 1:**
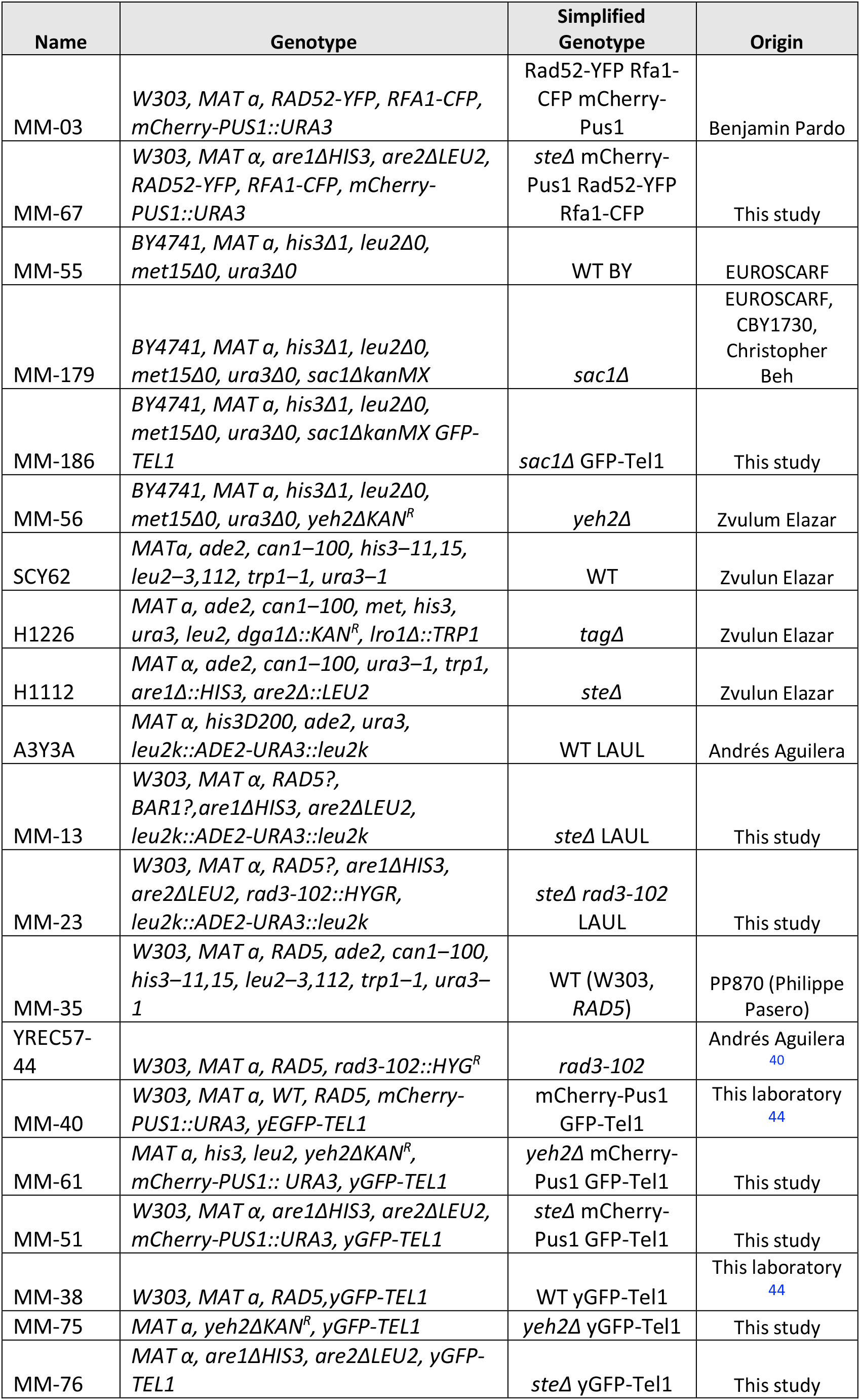
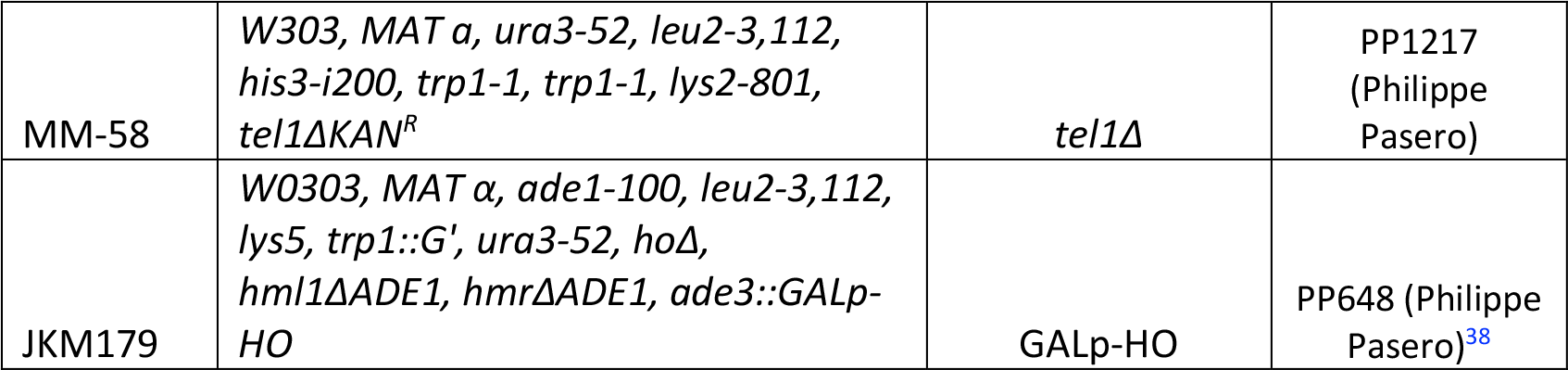
Strains used in this study

**Supplementary Table 2:**
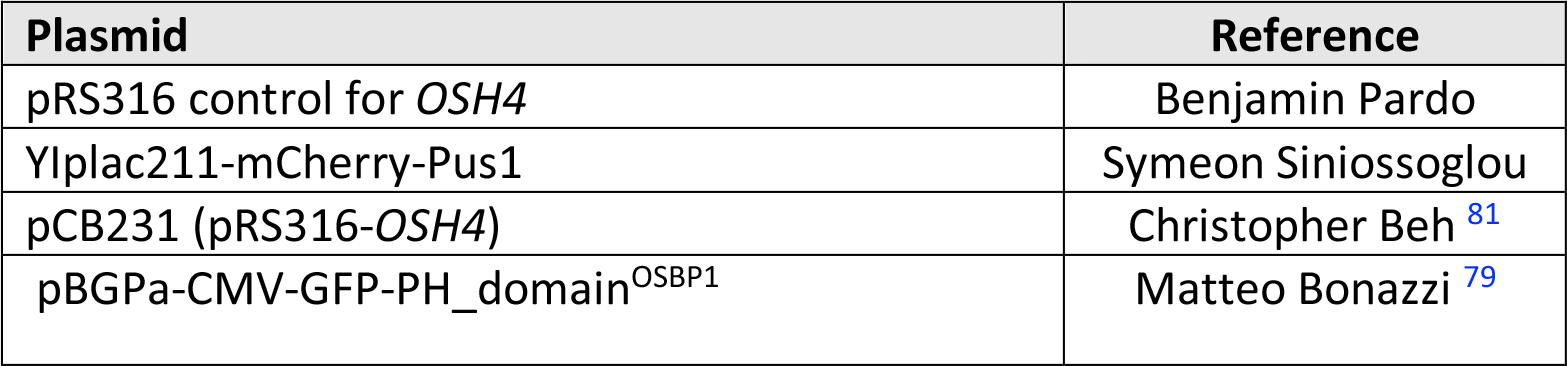
Plasmids used in this study

## Abbreviations

CPT: camptothecin
DDR: DNA Damage Response
DSB: DNA Double Strand Break
ER: endoplasmic reticulum
HGPS: Hutchinson-Gilford Progeria Syndrome
HR: homologous recombination
HU: hydroxyurea
LD: lipid droplets
MMS: methylmetane sulfonate
PI(4)P: phosphatidyl-inositol-4-phosphate
PLA: proximity ligation assay
ssDNA: single-stranded DNA
SEM: standard error of the mean
STEs: Sterol Esters
TAGs: TriAcylGlycerols
WT: wild type
5-FOA: 5-fluoro-orotic acid

## Acknowledgements

We are very grateful to Vincent Géli for the Rfa1-CFP-tagged strain; Corrado Santocanale for the anti-Rad53 antibody; Zvulun Elazar for *steΔ, tagΔ* and *yeh2Δ* strains; Katsunori Sugimoto for the *pGAL1p-FLAG-TEL1* vector; Michael Lisby for the original Rad52-YFP strain; Christopher Beh for the *sac1Δ* strain and the vector to overexpress Osh4; Bruno Mesmin and Bruno Antonny for the gift of schweinfurthin G; Symeon Siniossoglou for the YIplac211-mcherry-Pus1 vector; Andrés Aguilera for the *leu2k::ADE2-URA3::leu2k*-bearing and *rad3-102* strains; Jim Haber for the JKM179 strain to study resection; Matteo Bonazzi for the OSBP1-detecting biosensor and Urszula Hibner and Krzysztof Rogowski for the gift of Huh-7 and RPE-1 cells, respectively. We are very thankful to Jérôme Moreaux for supporting S.O. during the execution of this project. We acknowledge the imaging facility MRI, a member of the national infrastructure France-BioImaging supported by the French National Research Agency (ANR-10-INBS-04, Investissements d’avenir). SO was supported by a post-doctoral grant from La Ligue contre le Cancer. SK was supported by a SIRIC Montpellier Cancer Grant INCa_Inserm_DGOS_12553. AC and PP were supported by the MSDAvenir fund. The work in MM-C’s laboratory is supported by the ATIP-Avenir program, La Ligue contre le Cancer et l’Institut National du Cancer (PLBIO19-098 INCA_13832), France.

## Author contributions

Conceptualization, M.M.-C.; Methodology, S.O., S.K., C.S., O.S., B.P. and M.M.-C.; Investigation, S.O., S.K., C.S., O.S., J.A., B.P. and M.M.-C.; Writing – Original Draft, B.P. and M.M.-C.; Writing – Review & Editing, S.O., S.K., C.S., O.S., B.P., A.C., P.P. and M.M.-C.; Funding Acquisition, A.C., P.P. and M.M.-C.; Visualization, M.M.-C.; Supervision, M.M.-C.

